# “A feed-forward Ca^2+^-dependent mechanism boosting glycolysis and OXPHOS by activating Aralar-malate-aspartate shuttle, upon neuronal stimulation”

**DOI:** 10.1101/2021.02.02.429391

**Authors:** Irene Pérez-Liébana, Inés Juaristi, Paloma González-Sánchez, Luis González-Moreno, Eduardo Rial, Maša Podunavac, Armen Zakarian, Jordi Molgó, Beatriz Pardo, Jorgina Satrústegui, Araceli del Arco

**Affiliations:** Departamento de Biología Molecular, Centro de Biología Molecular Severo Ochoa, Consejo Superior de Investigaciones Científicas-Universidad Autónoma de Madrid (CSIC-UAM), Madrid, Spain. Instituto de Investigación Sanitaria Fundación Jiménez Díaz (IIS-FJD), Madrid, Spain; Facultad de Ciencias Ambientales y Bioquímica, Universidad de Castilla la Mancha, Toledo, Spain. Centro Regional de Investigaciones Biomédicas, Unidad Asociada de Biomedicina UCLM-CSIC, Toledo, Spain; Department of Structural and Chemical Biology, Centro de Investigaciones Biológicas Margarita Salas (CIB-CSIC), Madrid, Spain; Department of Chemistry and Biochemistry, University of California, Santa Barbara, CA 93106, USA; Université Paris-Saclay, CEA, Institut des Sciences du Vivant Frédéric Joliot, ERL CNRS n° 9004, Département Médicaments et Technologies pour la Santé (DMTS), Service d’Ingénierie Moléculaire pour la Santé (SIMoS), F-91191 Gif sur Yvette, France

**Keywords:** Neuronal metabolism, calcium regulation, glycolysis, cell respiration, malate aspartate shuttle, Aralar/AGC1/Slc25a12, mitochondrial calcium uniporter, genetically coded metabolite sensors

## Abstract

Calcium is an important second messenger regulating a bioenergetic response to the workloads triggered by neuronal activation. In cortical neurons using glucose as only fuel, activation by NMDA, which elicits a strong workload dependent on Na^+^ entry, stimulates glucose uptake, glycolysis, pyruvate and lactate production, and OXPHOS in a Ca^2+^-dependent way. We find that Ca^2+^-upregulation of glycolysis, pyruvate levels and respiration, but not glucose uptake, all depend on Aralar/AGC1/Slc25a12, the Ca^2+^regulated mitochondrial aspartate-glutamate carrier, component of the malate-aspartate shuttle (MAS). Ca^2+^-activation of MAS increases pyruvate production, which directly fuels workload-stimulated respiration. Also it stimulates glycolysis. MCU silencing had no effect indicating that none of these processes required mitochondrial Ca^2+^. The neuronal respiratory response to carbachol was also dependent on Aralar, but not on MCU. We also find that cortical neurons are endowed with a constitutive ER-to-mitochondria Ca^2+^ flow maintaining basal cell bioenergetics in which Ryanodine receptors, RyR2, rather than InsP_3_R, are responsible for Ca^2+^ release, and in which MCU does not participate. The results reveal that in neurons using glucose MCU does not participate in OXPHOS regulation under basal or stimulated conditions, while Aralar-MAS appears as the major Ca^2+^-dependent pathway tuning simultaneously glycolysis and OXPHOS to neuronal activation.

## Introduction

The mammalian brain is a prominent consumer of glucose and oxygen in the resting state. In the adult brain, neurons have the highest energy demand (Attwell & Laughlin, 2001), requiring continuous delivery of glucose from blood. Neuronal activation involves increases in cell workloads such as those caused by ion pumping (Attwell & Laughlin, 2001), and synaptic vesicle recycling (Rangaraju et al., 2014) in order to recover the basal state. In response to these workloads, neurons rapidly increase ATP production, mainly through increases in glycolysis and oxidative phosphorylation (OXPHOS) (Hall et al., 2012; Mergenthaler et al., 2013; Dienel, 2019; Rangaraju et al., 2014; Ashrafi and Ryan, 2017; Connolly et al., 2014). The mitochondrial response to workloads is immediate (Connolly et al., 2014; Llorente-Folch et al., 2013; 2015), and in the case of small workloads, it takes place without a detectable decrease in cytosolic ATP levels (Baeza-Lehnert et al., 2019), in dendrites (Gerkau et al., 2019) and the presynapse (Rangaraju et al., 2014).

It has been known for a long time that the rate of respiration depends not only on ATP demand but also on Ca^2+^ (Denton, 2009; Glancy and Balaban, 2012; Llorente-Folch et al., 2015), providing a feed-forward system to adjust OXPHOS to energy demand. This regulatory role of Ca^2+^ was shown to be independent of its effect in ATP demand (Llorente-Folch et al., 2013). The mechanism whereby this takes place is yet controversial. Indeed, Ca^2+^ may upregulate respiration thanks to its entry in mitochondria through the mitochondrial calcium uniporter complex (MCUC) (de Stefani et al., 2016), activating pyruvate dehydrogenase (PDH), phosphatase 1 (Pdp1) (Fecher et al., 2019), NAD^+^-linked isocitrate dehydrogenase (IDH3), and α-KGDH (Denton et al., 2009; Armstrong et al., 2014). These effects of Ca^2+^ would increase substrate supply to the respiratory chain and the rate of respiration in response to energy demand.

The finding that a global MCU deletion in outbred CD1 mice did not have any gross dysfunction (but was lethal in the inbred C57BL/6 strain) was at odds with the functions attributed to MCU in the control of bioenergetics (Pan et al., 2013; Wang et al., 2020). Very recently it has been shown that synapses using pyruvate and lactate rely on Ca^2+^-dependent mitochondrial ATP production (Ashrafi et al., 2020), thanks to an enrichment in MICU3 which lowers the threshold for Ca^2+^ uptake in neuronal mitochondria (Ashrafi et al., 2020). However, whether these mechanisms are actually relevant to control respiration in neurons, using glucose, awaits confirmation.

Another way in which Ca^2+^ may operate to increase coupled respiration is through an increase in the activity of mitochondrial metabolite carriers regulated by Ca^2+^, with Ca^2+^-binding motifs facing the intermembrane space (Satrustegui et al., 2007). For example, the aspartate-glutamate exchanger Aralar/AGC1/Slc25a12 is a component of the malate aspartate shuttle (MAS) which is regulated by cytosolic Ca^2+^, with half-maximal effects at around 300 nM (Pardo et al., 2006; Contreras et al., 2007). In brain mitochondria (Gelllerich et al., 2013) and neurons using glucose, Ca^2+^-activation of MAS drives pyruvate away from lactate and into mitochondria (Llorente-Folch et al., 2013).

SCaMC3/Slc25a23 is a Ca^2+^ regulated ATP-Mg/Pi exchanger present in brain, which also increases stimulated respiration response but only at very high workloads associated with the depletion of mitochondrial ATP, as occurs in response to PARP1 activation (Rueda et al., 2015).

In addition to its role during the response to workloads, Ca^2+^ has shown to be important in the regulation of basal respiration in many cell types (Cardenas et al., 2010; Mallilankaraman et al., 2012; Tomar et al., 2019; Filadi et al., 2018). A constitutive transfer of Ca^2+^ from ER-to-mitochondria was required to maintain basal respiration and ATP/AMP levels. It involved Inositol3P receptors (IP3R) on the ER, the MCU in mitochondria, and additional proteins involved in contacts between ER and mitochondria (Rossia et al., 2019). It is unknown if this pathway is also functional in neurons, and its specific components in ER and mitochondria, including MCU.

In this work, we have evaluated the contribution of these two systems, MCU and Aralar-MAS as Ca^2+^-regulation mechanisms to prime mitochondrial respiration in response to workloads and in the regulation of basal respiration in neurons using glucose as fuel. Glucose is the major neuronal fuel and any change in OXPHOS involves increased substrate supply to mitochondria that needs to be matched with increases in glycolysis. Therefore, a second fundamental question in this work is the influence of these Ca^2+^ regulation mechanisms in glycolysis itself.

We report that while *Mcu-*KD blocks the increases in mitochondrial Ca^2+^ elicited by carbachol (Cch) and NMDA, it is dispensable for regulating neuronal respiration using glucose in every condition studied, basal respiration, Cch-stimulated respiration and NMDA-stimulated respiration. We find a strong influence of calcium on the upregulation of respiration in response to different workloads as observed earlier, which is not due to the workload associated with Ca^2+^ removal (Llorente-Folch et al., 2013; Rueda et al., 2014). By employing new generation of florescent probes for cytosolic pyruvate, lactate, and glucose, we find that neurons respond to NMDA by a rapid increase in cytosolic pyruvate and lactate production, which is blocked in the absence of Ca^2+^. Remarkably, the Ca^2+^-dependent increase in pyruvate, but not lactate, is blunted in *Aralar*-KO neurons, revealing a dependence of Aralar-MAS in the provision of cytosolic pyruvate for mitochondrial respiration. Moreover, we show that NMDA triggers a Ca^2+^-dependent increase in glycolysis, which also depends on Aralar. Thus, in neurons using glucose, neuronal activity is met with a Ca^2+^-dependent feed-forward mechanism, which boosts simultaneously glycolysis and respiration by activating Aralar-MAS, as metabolic link between glycolysis and OXPHOS. In the absence of this single Ca^2+^ regulation mechanism, the Ca^2+^-dependent boost of pyruvate production and glycolysis is lost, and MCU does not compensate for this loss.

## Results

### Control of basal neuronal respiration by ER Calcium. The role of ER Ca^2+^ stores and MCU

In 2010, a new role for Ca^2+^ in the control of basal cell respiration was reported (Cardenas et al., 2010; Mallilankaraman et al., 2012). A constitutive transfer of Ca^2+^ from ER to mitochondria was shown to be required to maintain basal respiration and ATP/AMP levels, and to block macroautophagy *via* an AMPK-dependent pathway.

We have examined whether this constitutive Ca^2+^ transfer pathway plays a similar role in neurons, and the role of MCU in this process. As neurons express two types of Ca^2+^ release channels in the ER, IP3R and Ryanodine receptors (RyR) (Egorova and Bezprozvanny, 2018; Galeotti et al., 2008; Zalk et al., 2007) we have analyzed the role of both. To study the role of ER efflux *via* IP3R, we have employed a membrane permeable selective blocker of all types of IP3R, analog of Xestospongin B (XeB), a macrocyclic bis-1-oxaquinolizidine alkaloid originally extracted from the marine sponge *Xestospongia exigua* (Gafni et al., 1997; Jaimovich et al., 2005). This analog 3 ‘-desmethyl Xestospongin B (3-dmXeB) has been produced by chemical synthesis in the group of Prof. A. Zakarian and exerts similar action as XeB on IP3R.

Fig. 1 A shows 60 min preincubation of cortical neurons with 5 µM 3-dmXeB substantially reduces the Ca^2+^ induced by 200 µM (S)-3,5-dihydroxyphenylglycine (DHPG), an agonist of Group I Metabotropic glutamate receptors (Niswender and Conn, 2010) consistent with the ability of XeB to block IP3R (Jaimovich et al., 2005; Cardenas et al., 2010). 10 µM 3-dmXeB was also inhibitory but induced cellular damage and was not used any further. 3-dmXeB did not modify the basal respiration rate in the same cortical neurons, as determined with a Seahorse extracellular flux analyzer (Fig. 1 B) indicating that Ca^2+^ efflux from the ER through IP_3_R does not play a relevant role in the control of basal respiration.

**Figure 1.**
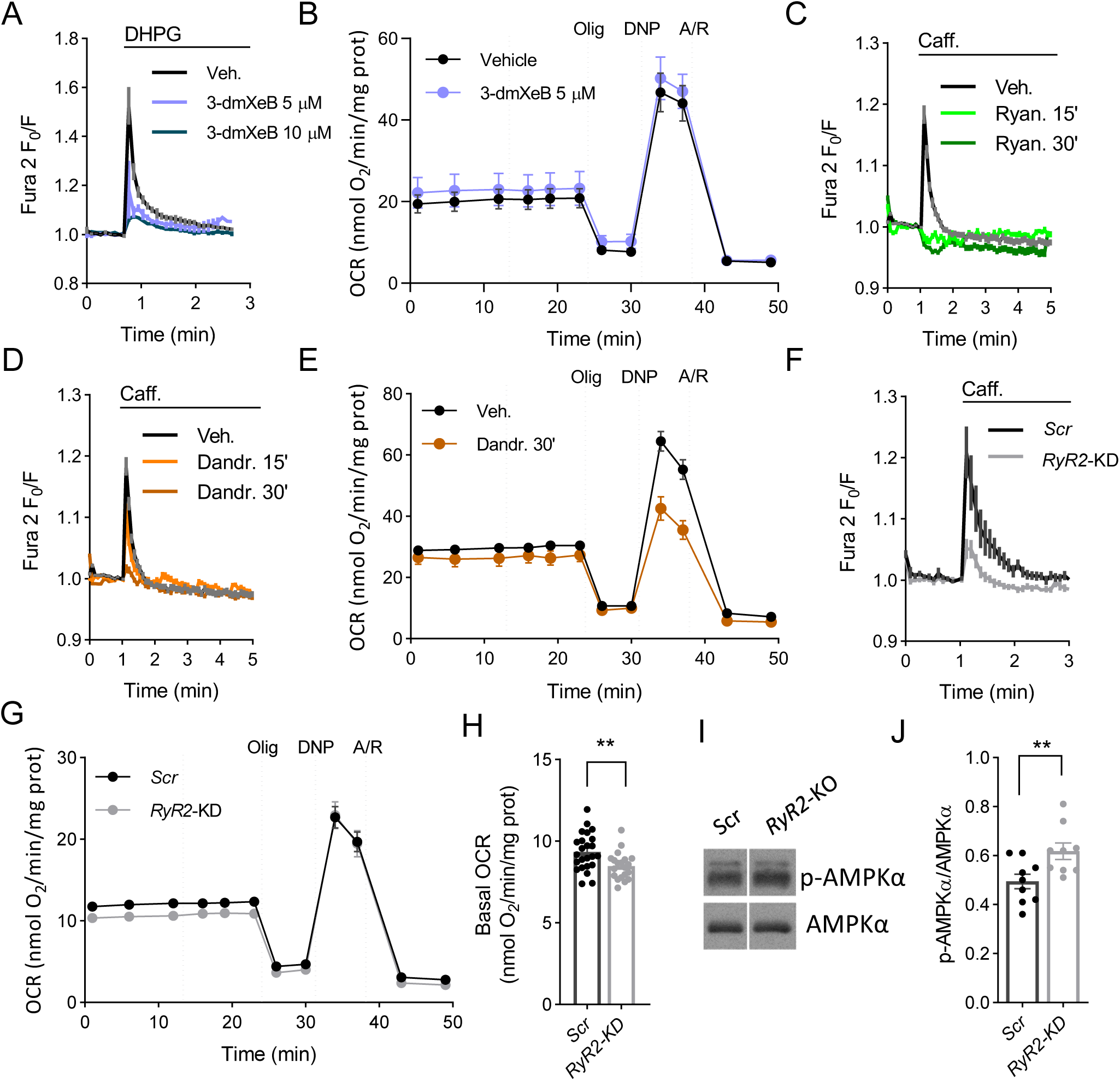
Ryanodine receptors control neuronal mitochondrial respiration. A: 5 and 10 µM 3-dmXeB substantially reduces the Ca^2+^ transients induced by 200 µM (S).-3,5-dihydroxyphenylglycine (DHPG), an agonist of Group I Metabotropic glutamate receptors in cortical neurons. Neuronal cultures were preincubated with 3-dmXeB 60 min before the experiment was started. B: 5 µM 3-dmXeB do not produce changes in basal respiration in cortical neurons. 3-dmXeB was added to the culture 60 min before the experiment was started. Mitochondrial function was determined through sequential addition of 6 μM oligomycin (Olig), 0.5 mM 2,4-dinitrophenol (DNP) and 1 μM/1 μM antimycin A/Rotenone (A/R) at the indicated time points. C-D: 15 mM Caffeine (Caff.) discharges ER calcium by activating RyR in cortical neurons at 9 DIV. Inhibition of the response to caffeine by preincubation with 50 µM ryanodine (Ryan.) (C) or 10 µM dandrolene (Dandr.) (D) during 15 or 30 min. E: Dandrolene caused a small decrease in basal OCR. Expressed as oxygen consumption rate (OCR; nmol O_2_/min/mg protein). Mean ± SEM, *n* = 9-11. F: *RyR2* silencing decreases caffeine induced Cai signal. G: *RyR2* silencing decreases basal OCR levels in neurons, representative experiment. Mean ± SEM, *n* = 7-8. H: Basal respiration without non-mitochondrial respiration; dots are individual data, bars are mean ± SEM, *n* = 14-16, \*\**p*-value = 0.0037, T-test. I: Representative WB analysis of p-AMPK and AMPK levels. J: Quantitative analysis from WB of p-AMPK/AMPK levels, dots are individual data, bars mean ± SEM, *n* = 9, ** *p*-value=0.0069 T-test.

In neurons, ER Ca^2+^ mobilization *via* activation of ryanodine receptors is involved in Calcium-induced Calcium release CICR (Shmigol et al., 1995; Solovyova et al., 2002; Patel et al., 2009). As observed in Fig. 1 C, 15 mM caffeine can discharge the ER Ca^2+^ store, as shown previously (Solovyova et al., 2002), resulting in an increase in cytosolic Ca^2+^. Preincubation with RyR inhibitors, such as Ryanodine (50 µM) (Solovyoka et al., 2002) (Fig. 1 C) or dandrolene (10 µM) (Liu et al., 2009; Oules et al., 2012) (Fig. 1 D) was able to block the release of Ca^2+^ caused by caffeine, confirming that these compounds block RyR in cortical neurons. Under these incubation conditions, the effect of ryanodine or dandrolene was restricted to the inhibition of caffeine-induced Ca^2+^ release, and not due to a discharge of ER Ca^2+^, since the size of this Ca^2+^ pool, as ascertained by the Ca^2+^ signal obtained by the administration of the Ca^2+^ ionophore Ionomycin, was unchanged (Fig. 1 Suppl. 1A). Interestingly, dandrolene also caused a small, but not significant, decrease in basal respiration (2.26 ± 0.10 *vs*. 2.07 ± 0.24, *n* = 11-9) indicative of a constitutive Ca^2+^ flow pathway from the ER to mitochondria involving RyR (Fig 1 E).

### Ryanodine receptors, but not MCU, are involved in the maintenance of basal respiration in cortical neurons

To verify the role of RyR in providing a constitutive Ca^2+^ flow regulating respiration in cortical neurons, we have silenced *RyR2*, the major RyR isoform in brain cortex and hippocampus (Galeotti et al., 2008; Zalk et al., 2007), using selected sequences used previously (Mu et al., 2014; More et al., 2018). *RyR2* silencing with two recombinant adeno-associated virus (rAAV)-mediated transduction of *RyR2*-directed small hairpin RNA (shRNA), or non-targeted control sequence (*Scrambled, Scr*) were verified by qPCR (Fig. 1 Suppl. 1 B) and resulted in a decrease in caffeine-induced Ca^2+^ release (Fig. 1 F). In addition, a significant decrease in basal respiration was observed in the *RyR2*-silenced neurons (*RyR2*-KD, Fig. 1 G-H), which affected both oligomycin-sensitive and -resistant respiration supporting a constitutive flow of ER Ca^2+^ to mitochondria involving, at least partially, RyR2 (Fig. 1 H).

Failure to maintain ER to mitochondria Ca^2+^ flux causes an increase in autophagy triggered by activation of AMPK (Cardenas et al., 2010; Mallilankaraman et al., 2012). Indeed, AMPK phosphorylation status increased in *RyR2*-KD neurons (Fig. 1 I, J). This was accompanied by an increase ECAR and a decrease in the OCR/ECAR ratio (Fig 1 suppl. 1C) suggesting a higher glycolytic flux in response to AMPK activation. In fact, when evaluating the adenine nucleotides levels present in these cells, *RyR2*-silencing did not lead to any significant changes (Fig. 1 Suppl. 1 D-H) in agreement with the robust energetic stability in neurons (Baeza-Lehnert et al., 2019).

We have next studied the involvement of the MCUC in this constitutive pathway. To this end, we silenced *Mcu*, the Ca^2+^ transporter component of MCU complex (de Stefani et al., 2016). Neurons were transduced with recombinant adeno-associated virus (rAAV) containing *Mcu*-directed shRNA, or non-targeted control sequence (*Scrambled, Scr*) (Qiu et al., 2013). Analysis by Western Blot shows a 64.5 ± 10.4 % decrease in MCU protein level (Fig. 2 A, B). Fig. 2 C shows that the basal FRET ratio of the mitochondrial Ca^2+^ probe 4mtD3cpv in *Mcu*-KD neurons is very similar to that of *Scr* neurons. Indeed, unchanged resting Ca^2+^-mit levels were observed for other tissue-specific MCU deletions, as muscle-(Gehrardi et al., 2018), cardiomyocyte-(Kwonk et al., 2015), or neuronal-specific *Mcu*-KD (Qiu et al., 2013; Ashrafi et al., 2020). Silencing of *Mcu*, had no effect on basal respiration with glucose, as the main substrate, when compared with neurons transduced with adenovirus carrying *Scrambled* sequences (Fig. 2 D-E). Similarly, neuron-specific knock down of MCU did not cause changes in basal respiration in cortical neurons (Nichols et al., 2018), and brown adipose tissue specific MCU ablation did not modify brown fat bioenergetics (Flicker et al., 2019). As MCU is clearly able to take-up Ca^2+^ in mitochondria (see below), we conclude that it is not required as part of the ER to mitochondria Ca^2+^ flow maintaining basal respiration in neurons.

**Figure 2.**
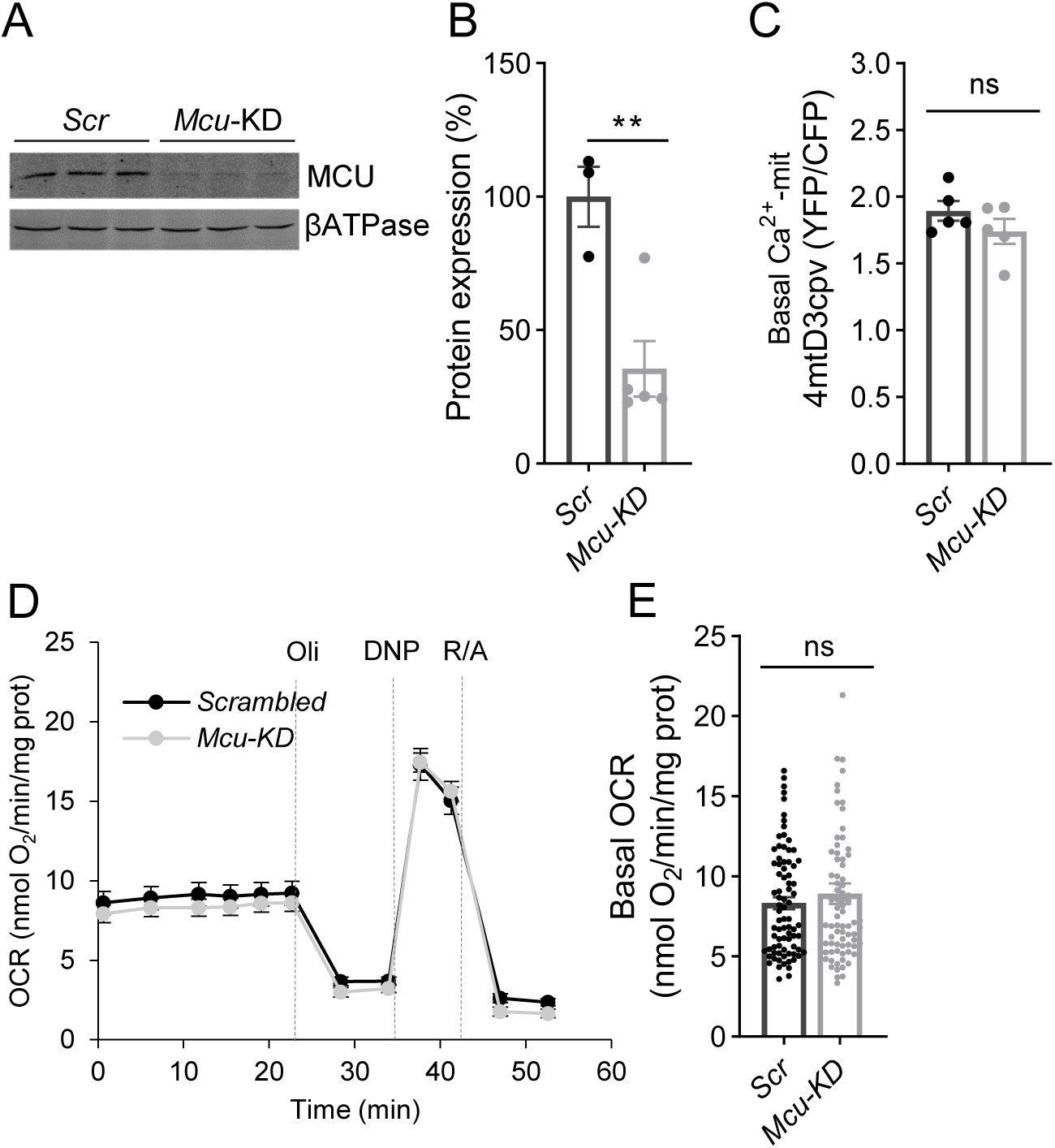
Basal mitochondrial Ca^2+^ and respiration do not change in *Mcu*-KD cortical neurons. A: Representative WB analysis of MCU levels in neurons transduced with rAAV containing MCU-directed (*Mcu*-KD) or *Scrambled* shRNA (*Scr*). βATPase was employed as charge control. B: Quantitative analysis from WB of MCU/ βATPase; dots are individual data, bars mean ± SEM, *n* = 3-5, \*\**p*-value = 0.0072, T-test. C: Basal levels of mitochondrial Ca^2+^ (Ca^2+^-mit) in *Scr* and *Mcu*-KD neurons transfected with 4mtD3cpv probe. Dots are individual data, bars mean ± SEM, *n =* 19-21 neurons per condition from 5 independent platings; *p*-value = 0.2335, T-test. D, E: Basal respiration in *Scrambled* and *Mcu*-KD neurons expressed as oxygen consumption rate (OCR; nmol O_2_/min/mg protein) mitochondrial function was determined through sequential addition of 6 μM oligomycin (Olig), 0.5 mM 2,4-dinitrophenol (DNP) and 1 μM/1 μM antimycin A/Rotenone (A/R) at the indicated time points. Dots are individual data, bars mean ± SEM, *n =* 26 per condition from 5 independent platings; *p*-value = 0.4193, T-test.

Therefore, our results show that a pathway for Ca^2+^ flow from ER to mitochondria that controls basal respiration exists in cortical neurons, but it is not dependent on IP3R on the ER or Ca^2+^ entry in mitochondria along MCU.

### The neuronal response to activation of acetylcholine receptors by carbachol

Carbachol (Cch) activation of acetylcholine receptors from cortical neurons resulted in a weak Ca^2+^-cyt signal that hardly reached mitochondria (Llorente-Folch et al., 2013). To study the impact of Ca^2+^ entry in mitochondria in response to Cch (250 µM) we have used mouse cortical neurons plated at high density and cultured for 8-9 days. These neurons showed spontaneous Ca^2+^ activity, consisting of repetitive Ca^2+^ transients occurring at the same time in all neurons (Fig. 3 A). These Ca^2+^ transients were blocked by treatment with the AMPA/Kainate receptor antagonist CNQX (10 μM), the NMDA receptor antagonist MK-801 (10 μM), or the Na^+^ channel blocker TTX (1 μM) (Fig. 3 A and results not shown), indicating that they depend upon a burst of action potentials (Opitz et al., 2002; Bacci et al., 1999; Young et al., 2005). Cch addition in the presence of TTX, which blocks other responses except those due to Ca^2+^ mobilization by IP3R, resulted in a weak response (Fig. 3 B) as reported previously (Delmas et al., 2002; Delmas et al., 2004). However, the most effective activation of IP3R is obtained when IP3 and Ca^2+^ are present together (Berridge, 1998; del Rio et al., 1999) and this explains the large effect of Cch in the absence of TTX, enhancing the spontaneous Ca^2+^ activity of the whole neuronal network, with an increase in the amplitude of the Ca^2+^ transients (Fig. 3 C, D).

**Figure 3.**
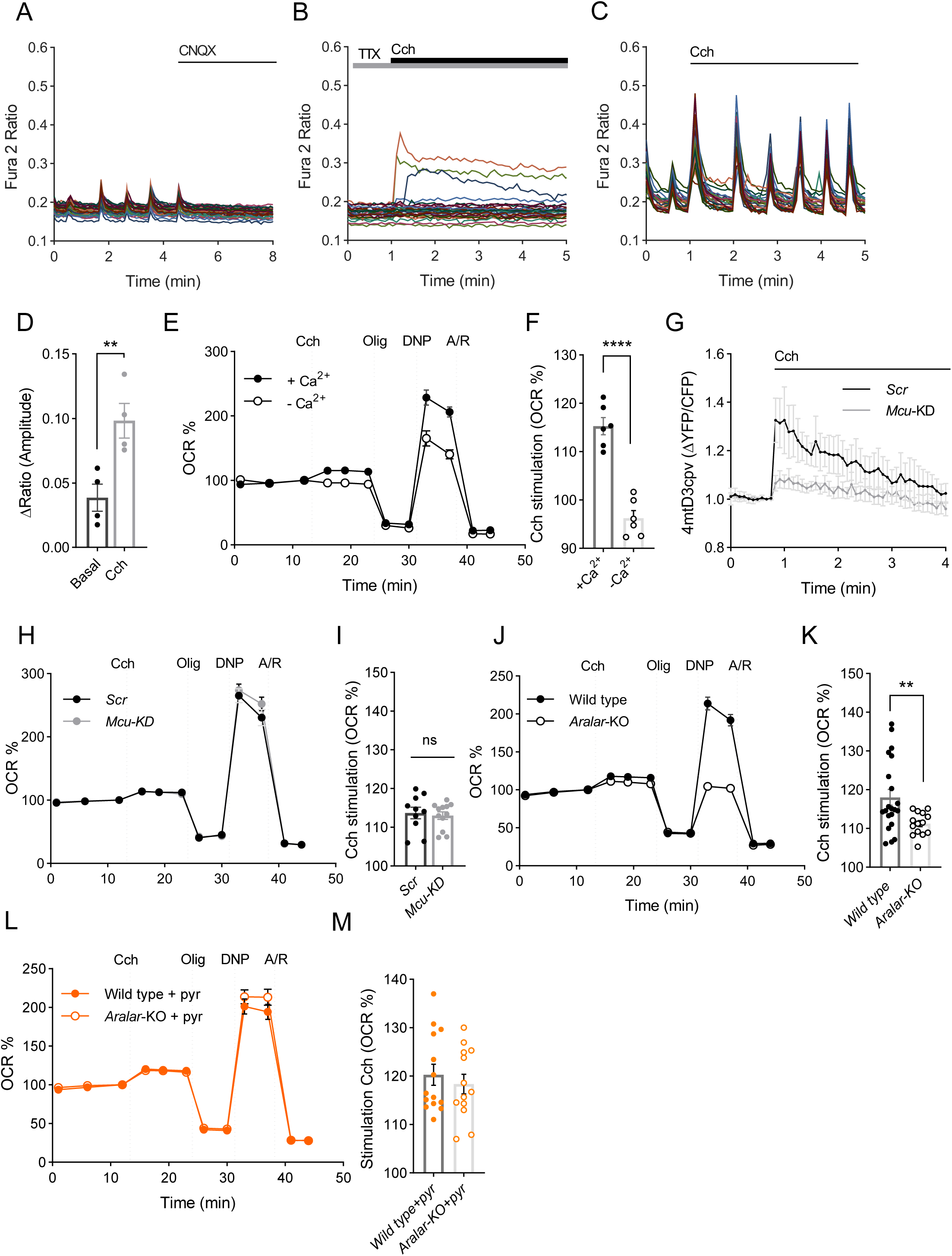
Cch stimulation of mitochondrial respiration depends on ARALAR-MAS pathway. A: Fura-2AM [Ca^2+^]i signals in neurons in HCSS medium containing 2 mM CaCl_2_ and 2.5 mM glucose, upon addition of 10 μM CNQX where indicated. B: Fura-2AM [Ca^2+^]i signals in cortical neurons in HCSS medium containing 2 mM CaCl_2_, 2.5 mM glucose and 1 μM TTX, upon addition of 250 μM Cch where indicated. C: Fura-2AM [Ca^2+^]i signals in neurons in HCSS medium containing 2 mM CaCl_2_, upon addition of 250 μM Cch where indicated. All graphs show a representative experiment; each trace corresponds to a single neuron from the same recording field. D: Quantification of peak amplitude as ΔRatio (F_340_/F_380_) ± SEM comparing basal spontaneous Ca^2+^ oscillations to Cch-enhanced Ca^2+^ oscillations. Data were obtained from 4 independent experiments. Means were compared using one-tailed T-test, *p*-value = 0.0064. E: Oxygen consumption rate expressed as percentage of basal OCR in WT neurons in medium containing 2 mM Ca^2+^ (+Ca^2+^) or in absence of Ca^2+^ (-Ca^2+^) showing the sequential injection of 250 μM Cch, and metabolic inhibitors: 6 μM Olig, 0.5 mM DNP and 1 μM/1 μM Ant/Rot at the indicated time points. F: Quantification of percentage of respiratory stimulation (OCR % 3 min after Cch addition) in WT neurons in presence or absence of Ca^2+^. Data were obtained from 6 independent experiments (*n* = 6). Means were compared using one-tailed T-test, *p* ≤ 0.0001. G: 4mt-D3cpv mitochondrial Ca^2+^ signals in *Scrambled* or *Mcu*-KD neurons upon addition of 250 μM Cch where indicated. Data are normalized to the initial values and are expressed as mean ± SEM. Data were obtained from 3 independent experiments (*n* = 8-11 cells). H: Oxygen consumption rate expressed as percentage of basal OCR in *Scrambled* and *Mcu*-KD neurons showing the sequential injection of 250 μM Cch, and metabolic inhibitors at the indicated time points. I: Oxygen consumption rate expressed as percentage of basal OCR in WT and *Aralar*-KO neurons showing the sequential injection of 250 μM Cch, and metabolic inhibitors: 6 μM Olig, 0.5 mM DNP and 1 μM/1 μM Ant/Rot at the indicated time points. F: Quantification of percentage of respiratory stimulation (OCR % 3 min after carbachol addition) in WT and *Aralar*-KO neurons. Data were obtained from 6 independent experiments (*n* = 21-14). Means were compared using one-tailed T-test, *p*-value = 0.0063. G: Oxygen consumption rate expressed as percentage of basal OCR in WT and *Aralar*-KO neurons in 2 mM pyruvate medium showing the sequential injection of 250 μM Cch, 6 μM Olig, 0.5 mM DNP and 1 μM/1 μM Ant/Rot at the indicated time points. H: Quantification of percentage of respiratory stimulation (OCR % 3 min after Cch addition) in WT and *Aralar*-KO neurons in 2 mM pyruvate medium. Data were obtained from 3 independent experiments (*n* = 11). Means were compared using one-tailed T-test, *p*-value = 0.53.

The robust Cch-induced increase in Ca^2+^-cyt transients results from IP3-dependent Ca^2+^ release from the ER and entry of extracellular Ca^2+^ (results not shown). The Cch-induced workload was fully Ca^2+^-dependent with no changes in Na^+^-cyt (Llorente-Folch et al., 2013) and accordingly, caused a stimulation of neuronal respiration only in the presence of extracellular Ca^2+^ (Fig. 3 E, F).

### Cch stimulation of mitochondrial respiration is independent of MCU pathway

To study the impact of MCU signaling on the Cch-induced increase in respiration, we silenced *Mcu*, and studied its effect on Cch-stimulated respiration. Down-regulation of MCU protein resulted in drastic block in mitochondrial Ca^2+^ signals, as Cch-driven mitochondrial Ca^2+^ rise was dramatically decreased in *Mcu*-silenced neurons (Fig. 3 G), while the increase in cytosolic Ca^2+^ transients evoked by Cch were the same in both *Scrambled* (*Scr*) or *Mcu*-KD neurons. No differences were found in amplitude or number of peaks after Cch addition (Fig. 3 Suppl. 1 A, B, C).

As described, basal respiration rates were similar in *Scrambled*- and *Mcu* silenced-neurons (Fig. 2 D, E). Strikingly, even though Cch-induced increase in matrix Ca^2+^ did not occur under *Mcu* silencing conditions, the stimulation of mitochondrial respiration by Cch was not affected, 113.57 ± 1.51 % in *Scrambled* and 113.02 ± 1.09 % (*p*-value = 0.71, T-test) in *Mcu*-silenced neurons over basal levels, respectively (Fig. 3 H, I).

### Cch stimulation of mitochondrial respiration depends on ARALAR-MAS pathway

Having shown that MCU is not required for Cch stimulation of respiration, we investigated the role of Aralar. We first verified that cytosolic Ca^2+^ signals induced by Cch in cortical neurons derived from wild type and *Aralar-*KO mice were the same (Cch enhanced Ca^2+^ oscillations to the same level; Fig. 3 Suppl. 1 D, E, F). Next, we studied Cch-stimulation of respiration in the two genotypes finding that Cch-stimulation of respiration is reduced by about 40 % in the absence of Aralar (Wild type neurons: 118.04 ± 1.03 % and *Aralar*-KO neurons: 111.2 ± 0.79 % increase over basal levels, *p*-value=0.0063, T-test) (Fig. 3 J, K). It is likely that the remaining stimulation of respiration is due exclusively to Ca^2+^-stimulated workload (i.e., Ca^2+^-induced ATP consumption), not to Ca^2+^ signaling. Although much smaller than that, dependent on Na^+^ entry, Ca^2+^ stimulated workload plays also a role in neuronal energy consumption (Llorente-Folch et al., 2013, Diaz-Garcia et al., 2020).

Activation by extramitochondrial Ca^2+^ of Aralar-MAS results in an increase in NADH production in neuronal mitochondria, which is blocked in *Aralar*-KO neurons, but reverted by exogenous pyruvate (Pardo et al., 2006). It has been proposed that Ca^2+^-activation of Aralar functions as a “gas pedal” to increase pyruvate formation (Gellerich et al., 2009; Gellerich et al., 2012; Gellerich et al., 2013). Indeed, Cch-stimulated respiration in *Aralar*-KO neurons was fully restored in the presence of 2 mM pyruvate (Fig. 3 L, M). Therefore, we conclude that the lack of Aralar results in a limitation in substrate (pyruvate) supply to mitochondria which prevents full Cch-induced stimulation of OCR.

### Role of mitochondrial calcium uniporter and Aralar-MAS in Ca^2+^-dependent stimulation of neuronal respiration triggered by NMDA

Activation of NMDA receptors leads to Na^+^ and Ca^2+^ entry in cortical neurons (Rueda et al., 2015). In order to restore the electrochemical gradients, Na^+^/K^+^-ATPases, plasma membrane Ca^2+^ ATPases (PCMA), and Na^+^/Ca^2+^ exchangers (NCX) are activated, leading to stimulated ATP use and to the upregulation of mitochondrial respiration to meet the energetic demands. In the absence of extracellular Ca^2+^, NMDA evokes a larger increase in neuronal Na^+^ than in its presence, which entails a larger workload in Ca^2+^-free than in Ca^2+^-containing medium (Rueda et al., 2015). In spite of this larger workload, Figure 4 A, B shows that acute addition of 25 µM NMDA to cortical neurons induced a smaller upregulation of OCR in the absence (-Ca^**2+**^, + 100 µM EGTA) than in the presence of Ca^2+^ (+Ca^**2+**^, 2 mM). This difference in the OCR response to NMDA resulted in a greater drop in cytosolic ATP levels in Ca^2+^-free than in Ca^2+^-containing medium (Rueda et al., 2015). These results clearly indicate an important role of Ca^2+^ upregulation of mitochondrial respiration in the neuronal response to NMDA.

**Figure 4.**
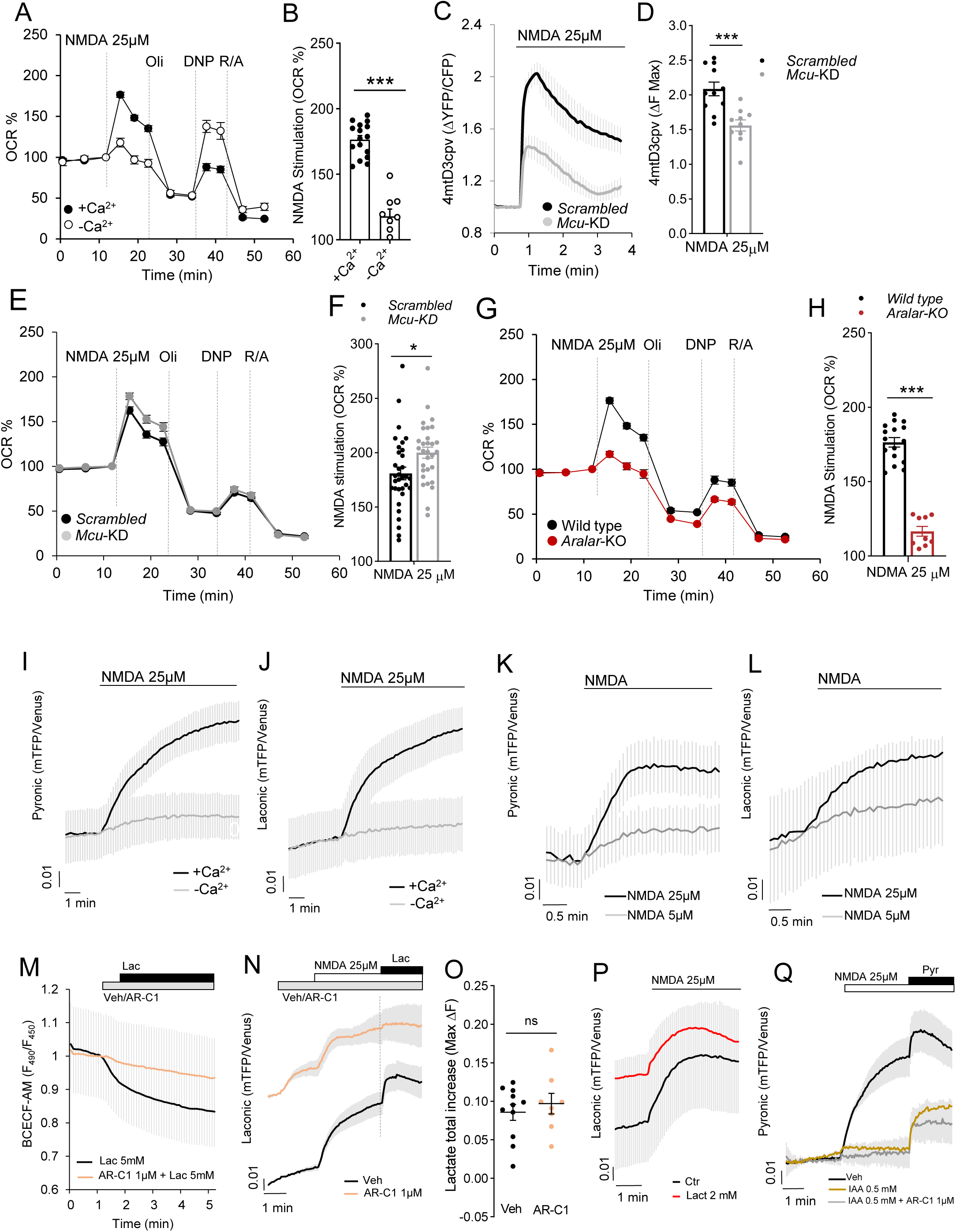
NMDA stimulation of OCR and formation of cytosolic pyruvate and lactate depends on Ca^2+^- and Aralar, but not on MCU. A: 25 μM NMDA-stimulated respiration in cortical neurons incubated in DMEM and 2.5 mM glucose, in the presence of 2 mM Ca^2+^ (+Ca^2+^) or without calcium (-Ca^2+^; plus 100 μM EGTA), expressed as percentage of basal values (OCR %). Mitochondrial function was determined through sequential addition of metabolic inhibitors: 6 μM Olig, 0.5 mM DNP and 1 μM/1 μM A/R at the indicated time points. B: Quantification of NMDA-stimulated OCR (%); mean ± SEM (bars), *n* = 9-16 (dots) from two different experiments; one-way ANOVA, \*\*\**p* ≤ 0.001, *post hoc* Bonferroni test. C: Changes in mitochondrial Ca^2+^ (Ca^2+^-mit) in *Scrambled* (*Scr*) and *Mcu*-silenced (*Mcu*-KD) neurons transfected with 4mtD3cpv probe, after 25 μM NMDA stimulation. D: Quantification of maximum Ca^2+^-mit increment (ΔF max) after NMDA stimulation. Data are mean ± SEM (bars), *n* = 10-11 (dots) from three independent experiments; T-test \*\*\**p*-value = 0.0006 E: 25 μM NMDA-stimulated respiration in *Scrambled* and *Mcu*-KD neurons, expressed as percentage of basal values (OCR %); F: Quantification of NMDA-stimulated OCR (%); mean ± SEM (bars), *n =* 30-33 (dots) from 5 independent experiments; T-test \**p*-value = 0.0173. G: 25 μM NMDA-stimulated respiration in wild type and *Aralar*-KO neurons, expressed as percentage of basal values (OCR %); H: Quantification of NMDA-stimulated OCR (%); mean ± SEM (bars), *n* = 9-16 (dots) from 2 independent experiments; T-test \*\*\*\**p*-value < 0.0001. I, J: Neurons transfected with Pyronic (I) and Laconic (J) probes to determine changes in cytosolic pyruvate (Pyr) and lactate (Lac), respectively, upon (+Ca^2+^) or absence (-Ca^2+^) of calcium. K, L: Dose-dependent effect of 5 μM and 25 μM NMDA induction of Pyr (K) and Lac (L) production. Data are mean ± SEM, *n* = 5-15 neurons per condition from 2-6 independent experiments. M. Intracellular pH variation determined in BCECF-AM-loaded neurons after 5 mM lactate (Lac) acute addition with or without 1 min pre-incubation with 1 μM MCT1/2 inhibitor AR-C155858 (AR-C1). Data are mean ± SEM, *n* = 6 per condition from 2 independent experiments. N: Lactate changes induced by 25 μM NMDA in neurons with a pre-addition of HCSS (Veh) of 1 μM AR-C1. O: Quantification of maximum fluorescence change (ΔF max) in Lac after Veh/1 μM AR-C1 and 25 μM NMDA. P: Lactate changes induced by 25 μM NMDA in neurons with extracellular 2 mM lactate (Lact) or in control (Ctr); Q: Pyruvate changes induced by 25 μM NMDA in neurons pre-incubated 30 min with HCSS (Veh), 0.5 mM iodoacetate (IAA) or IAA + 1 μM AR-C1 (IAA + AR-C1). Data are mean ± SEM, *n* = 8-20 per condition from 4-8 independent experiments for Pyronic and Laconic assays; 5 mM Pyr or Lac was added 3 min after NMDA as a control. The 4mtD3cpv, Pyronic, Laconic and BCECF-AM experiments were performed in HCSS, 2.5 mM glucose and 2 mM Ca^2+^ or 100 μM EGTA (-Ca^2+^).

To study the contribution of MCU and Aralar-MAS as Ca^2+^ regulation mechanisms in NMDA-stimulation of respiration, we started by silencing *Mcu* in cortical neurons, as described above. NMDA-stimulated increase in matrix Ca^2+^ was clearly smaller in *Mcu*-KD than *Scrambled*-neurons both with 25 µM NMDA (Fig. 4 C, D) and 5 µM NMDA (Fig. 4 suppl. 1 A, B). However, *Mcu*-silencing did not block completely NMDA-stimulated Ca^2+^ entry in mitochondria, a situation already described in cultured neurons (Qiu et al., 2013) and in agreement with the incomplete block of Ca^2+^-uptake in isolated brain mitochondria from global *Mcu*-KO mice (Hamilton et al., 2018; Szibor et al., 2020). This diminished Ca^2+^ transport into mitochondria was not translated into an increase in cytosolic Ca^2+^ in *Mcu*-silenced neurons; rather 5 and 25 μM NMDA caused a similar increase in cytosolic Ca^2+^ in *Mcu*-silenced and *Scrambled*-neurons (Fig. 4 suppl. 2 A-D).

Surprisingly, 5 (Fig. 4 suppl. 1 C, D) and 25 μM (Fig. 4 E, F) NMDA-stimulated OCR was not decreased by *Mcu-*silencing as expected if matrix Ca^2+^ played a role in stimulation of respiration; in fact the stimulation was slightly larger compared to *Scrambled*-neurons. The lack of inhibition of NMDA-stimulated respiration by *Mcu* silencing cannot be due to incomplete inhibition of Ca^2+^ entry in mitochondria, as that would result in small inhibition or no inhibition of NMDA-stimulated OCR, but not in an even larger response. In fact, *Mcu*-KD in cultured neurons was effective in preventing glutamate excitotoxicity (Qiu et al., 2013). Therefore, the results clearly indicate that cortical neurons using glucose do not require MCU and matrix Ca^2+^ to upregulate respiration, in response to NMDA-induced increase in workload. These results fully agree with the lack of effect of MCU deletion on respiration of brain or heart mitochondria on substrates using MAS, as shown by Szibor et al. (2020). To explore the role of Aralar we have studied the respiratory response to 5 and 25 µM NMDA. Stimulation of respiration by 5 (Fig. 4 Suppl. 1 E, F) and 25 µM (Fig. 4 G, H) NMDA was strikingly blunted in *Aralar*-KO as compared to wild type, as found in response to 50 μM glutamate (Llorente-Folch et al., 2016), while the absence of Aralar did not change the NMDA dependent rise in cytosolic and mitochondrial Ca^2+^ at the two NMDA concentrations (Fig. 4 Suppl. 2 E-L). Therefore, the presence of Aralar-MAS and not MCU has a critical role in Ca^2+^-regulation of respiration in response to NMDA.

### Activation of NMDA receptors increases cytosolic pyruvate and lactate production in a Ca^2+^– dependent way

The effect of the lack of Aralar on Cch-stimulated respiration is rescued by exogenous pyruvate (Fig. 3, L, M) (Llorente-Folch et al., 2013; 2016). Therefore, we have analyzed the impact of the presence or absence of Aralar in cytosolic pyruvate and lactate levels upon NMDA stimulation.

To this end, cortical neurons were transfected with Pyronic or Laconic FRET nanosensors (San Martín et al., 2014; San Martin et al., 2013) to register pyruvate or lactate levels, respectively as described previously (Juaristi et al., 2019a). The addition of 25 μM NMDA to neurons in the presence of Ca^2^ resulted in a prominent increase in both cytosolic pyruvate and lactate levels. The increase in both metabolites was drastically reduced in a Ca^2+^ free medium (Fig. 4 I, J). NMDA-induced pyruvate and lactate production was dependent on NMDA concentration (Fig. 4 K, L).

We next addressed the origin of lactate and pyruvate. Although purity of primary neuronal cultures is high, glial contamination can occur; and the surrounding astrocytes could provide lactate taken up by neurons. It has been shown that in a mixed culture, with physiological concentrations of glucose and lactate, the majority of neurons behaved as lactate consumers, and astrocytes as producers (Baeza-Lenhart et al., 2019).

In order to elucidate the origin of these metabolites, bilateral transport across plasma membrane was blocked through pharmacological inhibition of neuronal monocarboxylate transporter 2 (MCT2) with AR-C155858 (AR-C1), a potent inhibitor of both MCT1 and MCT2 (Ovens et al., 2010). MCT2 catalyzes the co-transport of a monocarboxylate (i.e., pyruvate, lactate, β-hydroxybutyrate) with a proton (Alvarez et al., 2003). To demonstrate the inhibitory effect of AR-C1 in our conditions, we have used the acidification associated with MCT2 activity as readout. Acute addition of 5 mM lactate (Lac) caused the expected intracellular acidification, as indicated from the drop of F_490_/F_450_ fluorescence ratio in BCECF-AM loaded neurons (Fig. 4 M). This was prevented by 1 min preincubation with 1 µM AR-C1 revealing the effectiveness of AR-C1 as MCT2 blocker. Acute inhibition of neuronal MCT2 with 1 µM AR-C1 leads to an accumulation of lactate *per se*, that further increases with 25 µM NMDA stimulation (Fig. 4 N); thus the total increase in cytosolic lactate after AR-C1 + NMDA is similar to that caused by NMDA alone (Fig. 4 N, O). These results show that lactate arises from neuronal synthesis rather than by capture from the external media, as shown in activated brain neurons (Díaz-García et al., 2017). It should be noted that neuronal cultures have a relatively large lactate production even in the basal state (Juaristi et al., 2019a, b), possibly larger than that of brain neurons, likely limited by astrocyte lactate buildup in the extracellular space. However, regardless of the actual basal state, NMDA clearly stimulates lactate production even in the presence of 2 mM lactate in the media (Fig. 4 P).

In the case of pyruvate, 1 µM AR-C1 also increased rather than decreased pyruvate accumulation (results not shown), indicating an internal origin of pyruvate. This origin is glycolytic, since 30 min pre-incubation with 0.5 mM iodoacetate (IAA), an inhibitor of glyceraldehyde-3-phosphate dehydrogenase (GAPDH), prevented neuronal generation of pyruvate (Fig. 4 Q), and partially inhibited lactate production (results not shown), confirming its glycolytic origin. Altogether, our results show that neurons stimulate glycolytic production of pyruvate and lactate upon 25 µM NMDA action in a Ca^2+^-dependent manner.

### NMDA-induced pyruvate, but not lactate, production drops in the absence of Aralar-MAS

NMDA (25 µM)-induced pyruvate production decreased in *Aralar*-KO neurons with the maximum pyruvate increase reduced by 44 % in comparison with wild type (Fig. 5 A, B; 0.059 ± 0.007 and 0.033 ± 0.006 for wild type and *Aralar*-KO). Pyruvate production in Ca^2+^-free media was largely abolished in *Aralar*-KO, and wild type neurons (Fig. 5 A-C). However, NMDA-induced lactate production was not affected by the absence of Aralar (Fig. 5 D), as reflected in the same maximum increase (Fig. 5 E; 0.093 ± 0.002 and 0.086 ± 0.002 for wild type and *Aralar*-KO), and in the same velocity of lactate production (Fig. 5 F; 0.063 ± 0.002 and 0.046 ± 0.001 for wild type and *Aralar*-KO). Ca^2+^ dependence of lactate production upon 25 µM NMDA stimulation was also observed in *Aralar*-KO neurons (Fig. 5 D-F).

**Figure 5.**
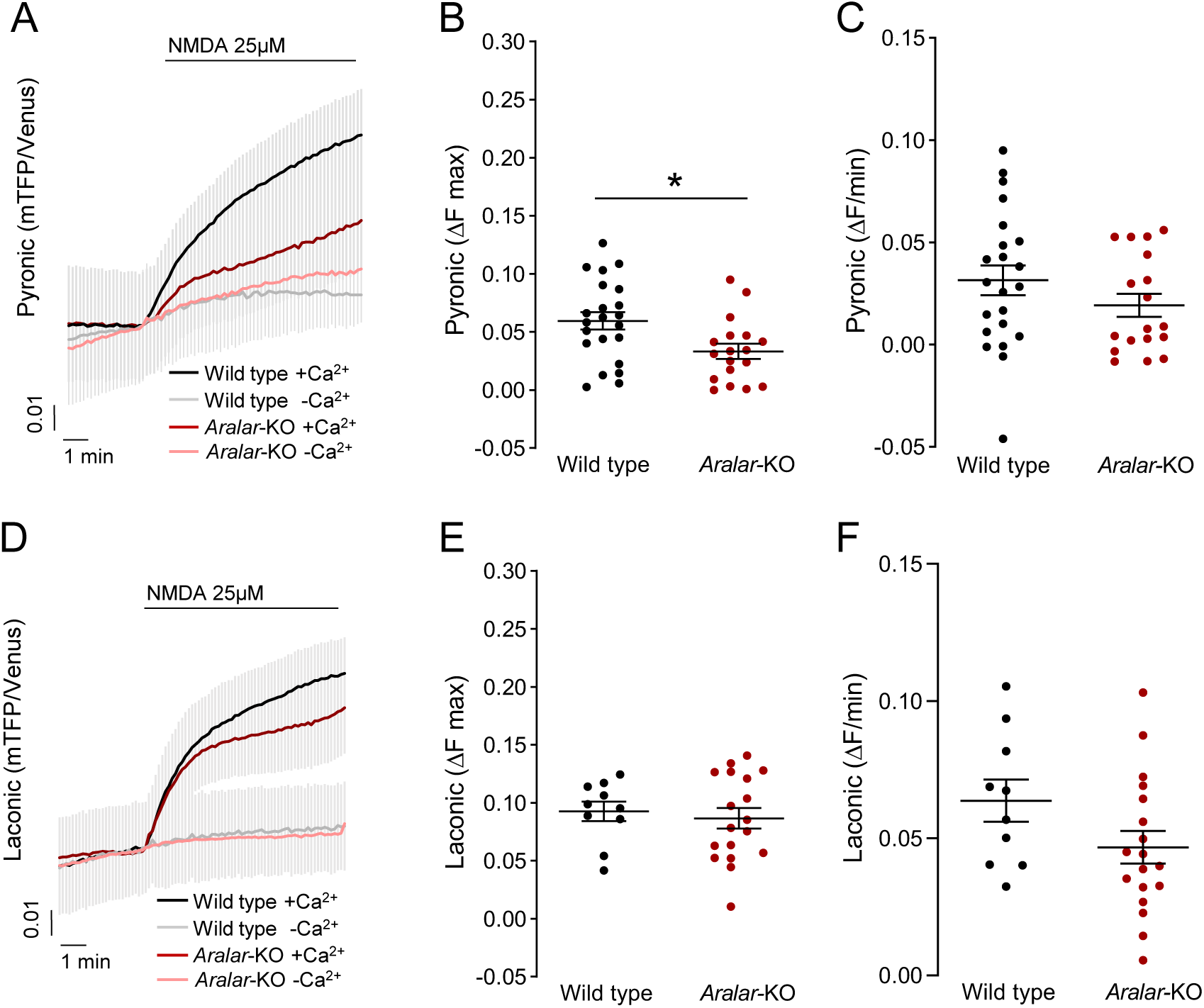
NMDA-induced changes in cytosolic pyruvate and lactate, in wild type and *Aralar*-KO primary neuronal cultures. A, D: Cortical neurons from wild type and *Aralar*-KO mice transfected with Pyronic (A) and Laconic (D) probes were stimulated with 25 μM NMDA in 2.5 mM glucose HCSS, in the presence of 2 mM Ca^2+^ (+Ca^2+^) or 100 μM EGTA (-Ca^2+^). FRET changes reporting cytosolic pyruvate or lactate levels are shown. B, E: Quantification of maximum fluorescence change (ΔF max) in Pyr (B) and Lac (E) after NMDA stimulation. C, F: Quantification of velocity of increase of Pyr (C) and Lac (F) as the increment of fluorescence ratio during the first 30 seconds after stimulation (ΔF/min). *n =* 10-22 per condition from 5-8 independent experiments for all Pyr and Lac assays; 5 mM Pyr or Lac was added 3 min after NMDA as a control. Data are mean ± SEM, T-test \**p* = 0.0133.

These results suggest that Aralar is required to stimulate mitochondrial respiration in response to an NMDA-induced workload in the presence of Ca^2+^ by providing pyruvate to mitochondria. The Ca^2+^-dependent stimulation of pyruvate generation is consistent with: (i) the stimulation of MAS by Ca^2+^ conferred by Aralar in isolated brain mitochondria (Pardo et al., 2006; Contreras et al., 2007), (ii) the stimulation by Ca^2+^ of respiration on glutamate + malate of brain mitochondria (Gellerich et al., 2012, 2013; 2009, Szibor et al, 2020), and (iii) the observed recovery of *Aralar*-KO deficient neuronal respiration by external pyruvate administration during stimulation (Fig. 3 L, M; Pardo et al., 2006; Llorente-Folch et al., 2013). On the other side, *Mcu*-KD had no influence on NMDA-induced production of pyruvate and lactate (Fig. 5 Suppl. 1 A-D).

### Activation of NMDA receptors activates glucose uptake and glycolysis in a Ca^2+^ dependent way

Having found that NMDA stimulation increases glycolytic pyruvate and lactate production in a Ca^2+^-dependent way, we wondered about the origin of glucose fueling this process and its stimulation by NMDA. To monitor cytosolic glucose, we have employed the glucose FRET nanosensor FLII^12^Pglu-700μδ6 (Takanaga et al., 2008) which is pH-independent, thanks to the replacement of eYFP by Citrine (Takanaga et al., 2008; Bittner et al., 2010). Indeed, NMDA induces a rapid cytosolic acidification (Fig. 6 A, see comparison with that caused by acute addition of HCl to BCECF-loaded neurons) as shown earlier (Rathje et al., 2013; Wu et al., 1999). However, the response to NMDA observed with FLII^12^Pglu-700μδ6 was an upward deflection of the FRET ratio consistent with an increase in glucose levels (Fig. 6 B). In contrast, HCl addition did not cause any increase in FRET ratio, but rather a gradual decrease probably due to side effects and probe quenching. Therefore, the FLII^12^Pglu-700μδ6 fluorescence changes observed in our experiments in primary cortical neurons are not due to acidification evoked by 25 μM NMDA, but really due to changes in cytosolic glucose levels. Neuronal stimulation is expected to decrease intracellular glucose levels, due to activation of glycolysis to cope with ATP demand. However, the results depicted in Fig. 6 C (black traces) show that FLII^12^Pglu-700μδ6 fluorescence ratio increases immediately after 25 μM NMDA stimulation, as observed previously in cerebellar granule neurons (Connolly et al., 2014). The rise in cytosolic glucose levels is maintained during a short period, and then steadily declines, suggesting an initial upregulation of glucose uptake or synthesis which is subsequently consumed.

**Figure 6.**
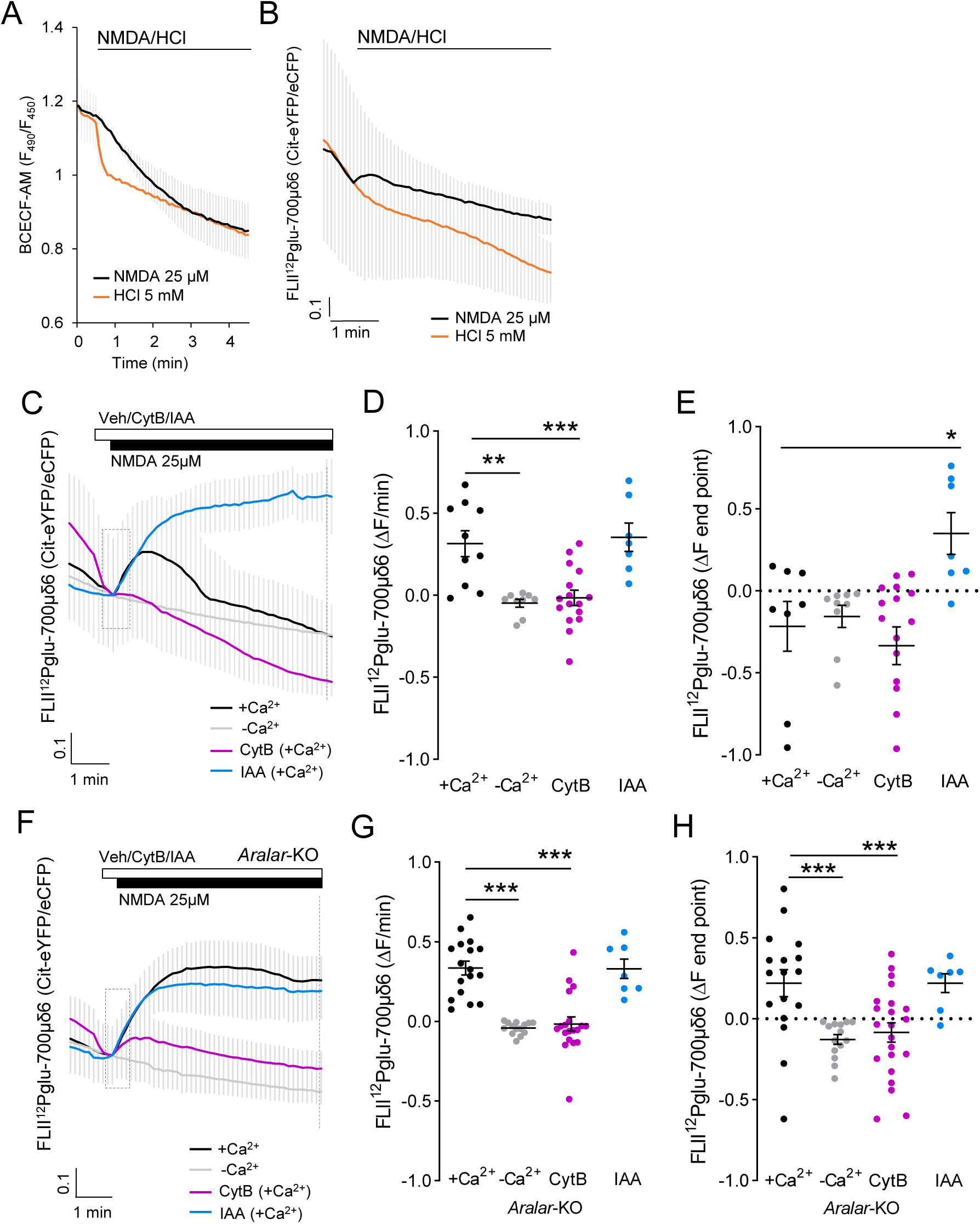
25 μM NMDA–induced changes in cytosolic glucose levels. A: Changes in cytosolic pH in BCECF-loaded neurons after 25 μM NMDA and 5 mM HCl acute addition. B: Changes in glucose levels in neurons transfected with FLII^12^Pglu-700μδ6 probe, after 25 μM NMDA and 5 mM HCl acute addition. Data are mean ± SEM, recordings from 5-6 cells per condition. C, F: 25 μM NMDA induced changes in glucose levels in wild type (C) and *Aralar*-KO (F) neurons, with or without calcium or with a 20 seconds pre-incubation with 50 μM cytochalasin B (CytB) or 0.5 mM iodoacetate (IAA). D, G: Velocity of increase of glucose levels during the first 30 seconds after stimulation (ΔF/min) in wild type (D) and *Aralar*-KO (G) neurons. E, H: FRET ratio change between pre-addition of NMDA and the end of the recording (ΔF end point) in wild type (G) and *Aralar*-KO (H) neurons. Mean ± SEM; recordings from 7-17 cells per condition from 3-5 independent experiments. One-way ANOVA, \**p* ≤ 0.05, \*\**p* ≤ 0.005, \*\*\**p* ≤ 0.001, *post hoc* Bonferroni test. ΔF at the end point after NMDA addition is higher in *Aralar*-KO than in wild type neurons; two-way ANOVA, \*\**p* ≤ 0.005, *post hoc* Bonferroni test. Assays using BCECF and FLII^12^Pglu-700μδ6 probes were performed in 2.5 mM glucose and 2 mM Ca^2+^ (+Ca^2+^) or 100 μM EGTA (-Ca^2+^) HCSS.

To determine if the increase in glucose obtained after NMDA exposure arises from uptake from the external medium, cortical neurons maintained in HCSS 2 mM Ca^2+^ and 2.5 mM glucose were treated with 50 µM cytochalasin B (CytB), an inhibitor of plasma membrane glucose transporters (Basketter and Widdas, 1978), 30 seconds prior 25 μM NMDA addition (Fig. 6 C, purple traces). NMDA-induced increases in Citrine-eYFP/eCFP fluorescence ratio in terms of slope (Fig. 6 D), and at the end of the experiment (ΔF end point; Fig. 6 E) are completely abolished in the presence of CytB, indicating that the increase in intracellular glucose arises from uptake from extracellular medium.

The following results, clarify that the subsequent steady decline in cytosolic glucose is due to glycolysis: a) NMDA-stimulated glucose consumption still continues in the presence of CytB (Fig. 6 C, purple traces); b) when 0.5 mM IAA was administered prior to NMDA addition (Fig. 6 C, blue traces), glucose levels dramatically increased consistent with a clear inhibition of glucose consumption by glycolysis. By employing mild electrical stimulation rather than bath application of NMDA, such an increase in cytosolic glucose (recorded with the same probe) and subsequent decrease was not observed (Baeza-Lehnert et al., 2019), possibly because increased uptake was balanced with increased glucose consumption.

Surprisingly, the NMDA-dependent rise in cytosolic glucose completely disappears in the absence of Ca^2+^, as shown by the block in the initial increase in glucose levels (Fig. 6 C, D, grey traces), and the NMDA-dependent decay in cytosolic glucose is also slower (ΔF end point; Fig. 6 E). Altogether, these results indicate that activation of NMDA receptors in cortical neurons upregulates glucose uptake from the external medium and its immediate consumption in glycolysis, both processes being dependent on Ca^2+^.

### NMDA-stimulation of glycolysis, but not glucose uptake, is strongly inhibited in *Aralar*-KO neurons

We next studied the involvement of Aralar in the upregulation of glucose uptake and glycolysis. Strikingly, NMDA-induced a much higher increase in glucose levels in *Aralar*-KO than in wild type neurons (Fig. 6 E and 6 H +Ca^2+^: 1.31 ± 0.05 *vs* 1.86 ± 0.02, in wild type and *Aralar*-KO; *p*-value = 0.0009, unpaired T-test), with no differences in velocity of increase between genotypes (Fig. 6 D and 6 G). NMDA-elevated glucose levels in *Aralar*-KO neurons were maintained throughout the recording interval instead of being immediately consumed as in wild type neurons (Fig. 6 F, H, black traces). NMDA-induced increase in glucose was blocked by incubation with 50 µM CytB (Fig. 6 F, G, purple traces) and in Ca^2+^-free medium (Fig. 6 F, grey traces), confirming that it was due to uptake from the external medium and was Ca^2+^-dependent also in *Aralar*-KO neurons. Thus, glucose uptake in neurons seems to be a Ca^2+^-dependent process, but independent from Aralar.

Inspection of Figure 6 shows differences in glucose consumption in wild type and *Aralar*-KO neurons. Unlike wild type neurons (Fig. 6 C), *Aralar*-KO neurons failed to consume cytosolic glucose in the absence or presence of CytB during the recording interval (Fig. 6 F, to compare black and purple traces). The very low cytosolic glucose consumption in *Aralar*-KO neurons was confirmed from the effect of incubation with 0.5 mM IAA. This inhibitor caused only a small increase in glucose accumulation (Fig. 6 H) when compared to its effects in wild type neurons (Fig. 6 E). Taken together, the results reveal a) that NMDA-induced glucose uptake is Ca^2+^-dependent in cortical neurons and is not affected by the lack of Aralar, and b) that NMDA-induced stimulation of glycolysis is also Ca^2+^-dependent and strictly requires Aralar.

Basal glucose consumption and lactate production are the same in wild type and *Aralar*-KO neurons (0.40 ± 0.08 and 0.43 ± 0.08 μmol glucose/mg/h, and 0.42 ± 0.05 and 0.44 ± 0.07 μmol lactate/mg/h respectively) (Juaristi et al., 2017). In the presence of Ca^2+^, NMDA-induced stimulation of glycolysis is much smaller in *Aralar*-KO than in wild type neurons, both in the absence (compare the glucose disappearance rate in WT and *Aralar*-KO in Figs. 6 C and 6 F, black traces) or presence of cytochalasin B (compare purple traces in Fig. 6 C and Fig 6 F). This is followed by a smaller stimulation of pyruvate formation (Fig. 5 A) and respiration (Fig. 4 G, H) while lactate formation was not significantly altered (Fig. 5 D). This entails that the fate of the small NMDA-increase in glycolysis in *Aralar*-KO neurons is largely lactate, with a diminished pyruvate formation and respiration.

## Discussion

### Constitutive Ca^2+^ flow from ER to mitochondria maintains basal respiration in neurons, depends on RyR2 and not IP3R and does not require MCU

The constitutive Ca^2+^ flow from ER to mitochondria has been shown to regulate basal respiration in many cell lines (Cardenas et al., 2010; Filadi et al., 2018), hepatocytes (Tomar et al., 2019) and skeletal muscle (Diaz-Vegas et al., 2018). We find that such a pathway also operates in neurons, with RyR2, but not IP3R, as source of ER Ca^2+^. This would seem surprising since IP3R are in close apposition to mitochondria, and IP3R1 is abundant in brain neurons, particularly in Purkinje neurons (Egorova and Bezprozvanny, 2018). However, the role RyR in controlling basal respiration is not new; RyR and IP3R control basal respiration in skeletal muscle (Diaz-Vegas et al., 2018). In fact, RyR are also close to mitochondria in neurons (Jakob et al., 2014) facilitate a rapid Ca^2+^ transfer from ER/SR to the matrix (Hajnóczky and Csordás, 2002), participate in MAM formation in brain (Volgyi et al., 2018; Ma et al., 2017) and are positioned in MCU enriched nanodomains in heart mitochondria (de la Fuente et al., 2018). Therefore, in an adequate position to participate in a constitutive Ca^2+^ flow to mitochondria.

We find that *Mcu*-KD has no influence on neuronal basal respiration. It could be argued that this is because of compensatory adaptations to MCU deletion masking its true role. For example, a well-known function of MCU, mitochondrial Ca^2+^ overload and mPTP opening, which is involved brain hypoxia/ischemia injury (Schinzel et al., 2005), remained unchanged in global *Mcu*-KO mice (Nichols et al., 2017). However, it was rescued in mice with tamoxifen-induced neuron-specific MCU ablation (Nichols et al., 2018), suggesting compensatory adaptations in the global, but not neuron-specific KO (Nichols et al., 2018, Bas-Orth et al., 2019). By using shRNAs to acutely silence *Mcu* in neurons, our approach is not expected to cause compensatory effects and therefore, the lack of effect of *Mcu*-KD in basal respiration (this work and Nichols et al., 2018) shows that MCU is not part of the ER to mitochondria Ca^2+^ flow regulating basal energetics in neurons using glucose. The Ca^2+^ sensor on mitochondria remains to be identified.

### Stimulation of mitochondrial respiration by agonists. Stimulation by Carbachol requires Aralar-MAS but not MCU

Our results clearly show that in cortical neurons with spontaneous Ca^2+^ activity Cch induces an increase in Ca^2+^ oscillations and an increase in neuronal respiration. Cch-induced Ca^2+^ oscillations reached mitochondria resulting in an increase in matrix Ca^2+^. Thus, matrix Ca^2+^ and activation of mitochondrial dehydrogenases (Glancy and Balaban, 2012) was a possible mechanism explaining the stimulation of respiration. Surprisingly, although matrix Ca^2+^ signals were definitely reduced in *Mcu* silenced neurons, *Mcu*-KD did not affect Cch-stimulated respiration.

In contrast, Cch-stimulation of respiration was blunted in *Aralar*-KO neurons. This reflects a limitation in substrate supply to mitochondria, as pyruvate addition abolished the differences in Cch-stimulated OCR between *Aralar*-KO and WT neurons. As neither MCU nor Aralar deficiency altered Cch-induced cytosolic Ca^2+^ oscillations, these results suggest that the Aralar-MAS pathway, but not MCU-driven Ca^2+^ entry in mitochondria is involved in boosting respiration by Cch-enhanced Ca^2+^ oscillations. The mechanism whereby Aralar-MAS regulates respiration in response to Cch becomes apparent from the analysis of the neuronal response to NMDA.

### Stimulation of mitochondrial respiration by agonists. NMDA-stimulation in cortical neurons causes Ca^2+^ dependent increases in glucose uptake, glycolysis, pyruvate and lactate formation and respiration

We have shown that NMDA exposure results in a Ca^2+^-dependent stimulation of respiration, and Ca^2+^-dependent increases in cytosolic pyruvate and lactate, both of which arose from glycolysis, and not from uptake from the external medium. It should be noted that following the application of a smaller workload, short theta bursts (40 pulses in 11 seconds) changes in lactate and pyruvate were not detected in hippocampal neurons (Baeza-Lenhart et al., 2019) probably because of the small dynamic range of the probe, especially around basal metabolite levels (San Martin et al., 2014, Juaristi et al., 2019a). This may explain the variability in stimulation-induced increase in Pyronic signal in WT and *Aralar*-KO cultures found in this study. However, in brain neurons increases in lactate and pyruvate have been observed after electrical stimulation (Díaz-García et al., 2017; Díaz-García and Yellen, 2019, Diaz-Garcia et al., 2020), or by arousal-evoked cortical activity (Zuend et al., 2019).

NMDA also triggers a Ca^2+^-dependent disappearance of cytosolic glucose blocked by inhibitors of glycolysis, showing that neurons respond to NMDA with a Ca^2+^-dependent stimulation of glycolytic glucose utilization. Additionally, NMDA stimulates Ca^2+^-dependent glucose uptake.

The power of the Ca^2+^ regulation mechanism(s) becomes apparent by comparison with the difference in workload between the conditions where Ca^2+^ is absent or present, in other words, the difference in ATP demand between these two conditions. The major contributor to neuronal workload is Na^+^ entry (Attwell and Laughlin, 2001, Baeza-Lehnert et al., 2019) and this is greatly reduced when external Ca^2+^ is present (Llorente-Folch et al., 2013; Rueda et al., 2015). Thus, the large NMDA-dependent stimulation of respiration, provision of pyruvate and lactate and glycolytic flux deployed in the presence of Ca^2+^ all take place in the face of a lower workload than in the absence of Ca^2+^. In the absence of Ca^2+^, the larger workload meets with non-operative Ca^2+^-regulation mechanism(s) and cytosolic ATP levels fall precipitously (Rueda et al., 2015).

It makes sense that Ca^2+^ regulates at the same time respiration, substrate supply to mitochondria, glycolysis and glucose uptake in response to an increase in workload. As neurotransmission is linked to Ca^2+^ signals, Ca^2+^ regulation of all these steps can be taken as a feed-forward mechanism to increase glycolysis and mitochondrial function upon initiation of the workload itself. But does this occur through a single mechanism or are there several Ca^2+^ dependent individual steps controlling the fate of glucose in neurons up to its combustion in mitochondria? Based on our results we suggest two Ca^2+^ dependent steps, one regarding glucose uptake and another one Aralar-MAS driven glycolysis and OXPHOS regulation. Our results show that glucose uptake has a specific Ca^2+^ regulation system independent of glycolysis. An increase in glucose uptake upon synaptic activity or NMDA-stimulation has been observed in both cerebellar and cortical neuronal cultures (Connolly et al., 2014, Ferreira et al., 2011). It required GLUT3/Slc2a3 translocation to the plasma membrane, via activation of AMPK (Connolly et al., 2014) or nNOS and cGMP-dependent protein kinase (Ferreira et al., 2011). However, the whole process is slower (10-15 min, Ferreira et al., 2011) than the rapid Ca^2+^-dependent increase in glucose uptake observed in this study. The major glucose transporter in muscle, GLUT4/Slc2a4, is translocated to the plasma membrane via a Ca^2+^-dependent pathway involving CaMKKβ (calmodulin-dependent protein kinase kinase *beta*) or a different CaM kinase and AMPK (Green et al., 2011; Kim et al., 2019; Angin et al., 2014). Ashrafi and Ryan (2017) have found that, similarly to muscle, activity at synapses triggers the rapid (seconds) translocation of GLUT4 to the plasma membrane, driven by AMPK. As CaMKKβ is involved in AMPK activation by depolarization in neurons (Kawashima et al., 2014), it is possible that this mechanism provides Ca^2+^ sensitivity to the NMDA-dependent increase in glucose uptake found in this study.

Regarding the other steps beyond glucose uptake, in one possible scenario there may be at least two independent Ca^2+^ regulated ones. One is Ca^2+^ regulation of glycolysis, and the other is Ca^2+^ activation of mitochondrial dehydrogenases (Denton, 2009) which may boost the activity of the Tricarboxylic Acid Cycle, and the use of pyruvate when Ca^2+^ enters the matrix. Ca^2+^-stimulation of glycolysis may involve a Ca^2+^-dependent activation of PFKFB3 (Rider et al., 2004). PFKFB3 is activated by AMPK (Almeida et al., 2004), which in turn is phosphorylated by CAMKKβ in response to rise in cytosolic Ca^2+^ (Sanders et al., 2007; Xiao et al., 2011). However, expression of PFKFB3 in neurons is low due to continuous degradation (Herrero-Méndez et al., 2009), and the Yellen group has shown that AMPK is not involved in regulation of glycolysis in stimulated neurons (Díaz-García et al., 2020) suggesting that this mechanism does not operate in response to acute NMDA stimulation. On the other hand, Ca^2+^ activation of mitochondrial dehydrogenases has been taken as the main feed-forward mechanism in which Ca^2+^ increases OXPHOS in response to workloads (Territo et al., 2000; Flicker et al., 2019) and requires MCU.

In another scenario, glycolysis and OXPHOS are Ca^2+^-modulated by the same mechanism, Aralar-MAS. The results from this study support this second possibility.

### Ca^2+^-dependence of NMDA-stimulation of glycolysis, pyruvate formation and respiration depends on Aralar-MAS and not MCU

Glycolysis can only proceed if there is sufficient supply of NAD^+^. The oxidation of glucose requires NAD^+^ (2 molecules for each glucose) that become NADH in the glyceraldehyde 3-phosphate dehydrogenase (GAPDH) reaction. Therefore, to sustain glycolysis, NAD^+^ needs to be continuously regenerated either by lactate dehydrogenase with conversion of pyruvate to lactate, or by NADH shuttle systems of which MAS is the prevalent in neurons (Juaristi et al., 2019a). MAS diverts pyruvate away from lactate and into OXPHOS and is regulated by cytosolic Ca^2+^ (Pardo et al., 2006; Contreras et al., 2007; Llorente-Folch et al., 2013; Rueda et al., 2014; Gellerich et al., 2012, 2013; Llorente-Folch et al., 2015), providing a potential feed-forward mechanism linking early Ca^2+^ signals arising from synaptic activity to both glycolysis and OXPHOS, as suggested earlier (Ashrafi and Ryan, 2017).

The results from the present study indicate that in neurons using glucose NMDA triggers a substantial workload met by a Ca^2+^-regulated response in which Ca^2+^ boosts glycolysis and OXPHOS by means of Ca^2+^ upregulation of Aralar-MAS. Indeed, Aralar deletion prevents NMDA effects on Ca^2+^-stimulation of glycolysis, pyruvate formation and respiration. In contrast *Mcu*-KD neurons were unaffected in terms of NMDA-stimulation of respiration or pyruvate production. This indicates that MCU-induced Ca^2+^ entry in mitochondria and activation of mitochondrial dehydrogenases (McCormack and Denton, 1993) is dispensable in the stimulation of respiration of neurons using glucose as also clearly shown by Diaz-Garcia et al., (2020). In other words, it entails that cytosolic Ca^2+^ regulation of Aralar-MAS, by increasing glycolysis and pyruvate levels, is sufficient to increase pyruvate transport in mitochondria and respiration. This agrees with the finding that matrix Ca^2+^-dependent PDH dephosphorylation has a rather limited role in controlling the production or uptake of cytosolic pyruvate and mitochondrial respiration in isolated mitochondria and neurons using glucose (Szibor et al., 2020, Llorente-Folch et al., 2015), and with the recent findings of Ashrafi et al., (2020), showing that the supply of ATP for synaptic vesicle endocytosis with glucose as fuel does not depend on MCU.

The critical role of increasing cytosolic pyruvate in regulating respiration may be associated with the low affinity for pyruvate of the mitochondrial pyruvate carrier, MPC, with a Km of about 0.15 mM (Halestrap, 1975) a concentration much higher than cytosolic pyruvate levels in astrocytes and neurons using glucose (San Martin et al., 2014; Baeza-Lenhert al., 2019, Juaristi et al., 2019a, this study).

Finally, it is important to emphasize that the reliance on Aralar-MAS as Ca^2+^ regulation switch connecting neuronal activity to metabolic response rather than MCU is limited to the use of glucose as substrate. Using a mixture of lactate and pyruvate instead of glucose, Ashrafi et al., (2020) have found that MCU is essential for synaptic vesicle endocytosis and the maintenance of ATP homeostasis after electrical stimulation in nerve terminals. By providing external pyruvate and not glucose, Aralar-MAS function is bypassed allowing a role of MCU and possibly matrix Ca^2+^-activation of OGDH IDH-NAD and PDH in the control of respiration. It is likely that matrix pyruvate levels need to increase beyond those present in neurons using glucose to allow a functional role of Ca^2+^-dependent activation of mitochondrial dehydrogenases in upregulating neuronal respiration. Further studies will be needed to clarify this point.

## Acknowledgments

This work has been funded by grants from the Spanish Ministry of Science, Innovation and Universities SAF2014-56929R (to JS and BP), SAF2017-82560-R (to AdelA and BP), by a grant from Fundación Ramón Areces (to JS), and by an institutional grant from the Fundación Ramón Areces to the Centro de Biología Molecular Severo Ochoa (CBMSO). I P-L and L G-M were recipients of predoctoral fellowships from MINECO and P G-S was the recipient of a postdoctoral research contract from Comunidad de Madrid. The authors thank Isabel Manso, Beatriz García and Bárbara Sesé for technical support and the unit of Optical and Confocal Microscopy from CBMSO. The qPCR experimental development was provided by the Genomics and NGS Core Facility at the CBMSO.

## Author contributions

IP-L, IJ and PG-S performed and analyzed the experiments, L G-M performed and analyzed AMPK WB and qPCR assays and contributed to experiments using viral vectors. MP, AZ, and JM, contributed providing reagents. ER, performed ATP determinations. BP, contributed to animals and neuronal cultures supervision. IP-L, IJ, PG-S and JS wrote the manuscript. AdA and JS conceived and supervised the whole project and supervised the writing of the manuscript. All authors edited and approved the manuscript.

## KEY RESOURCES TABLE

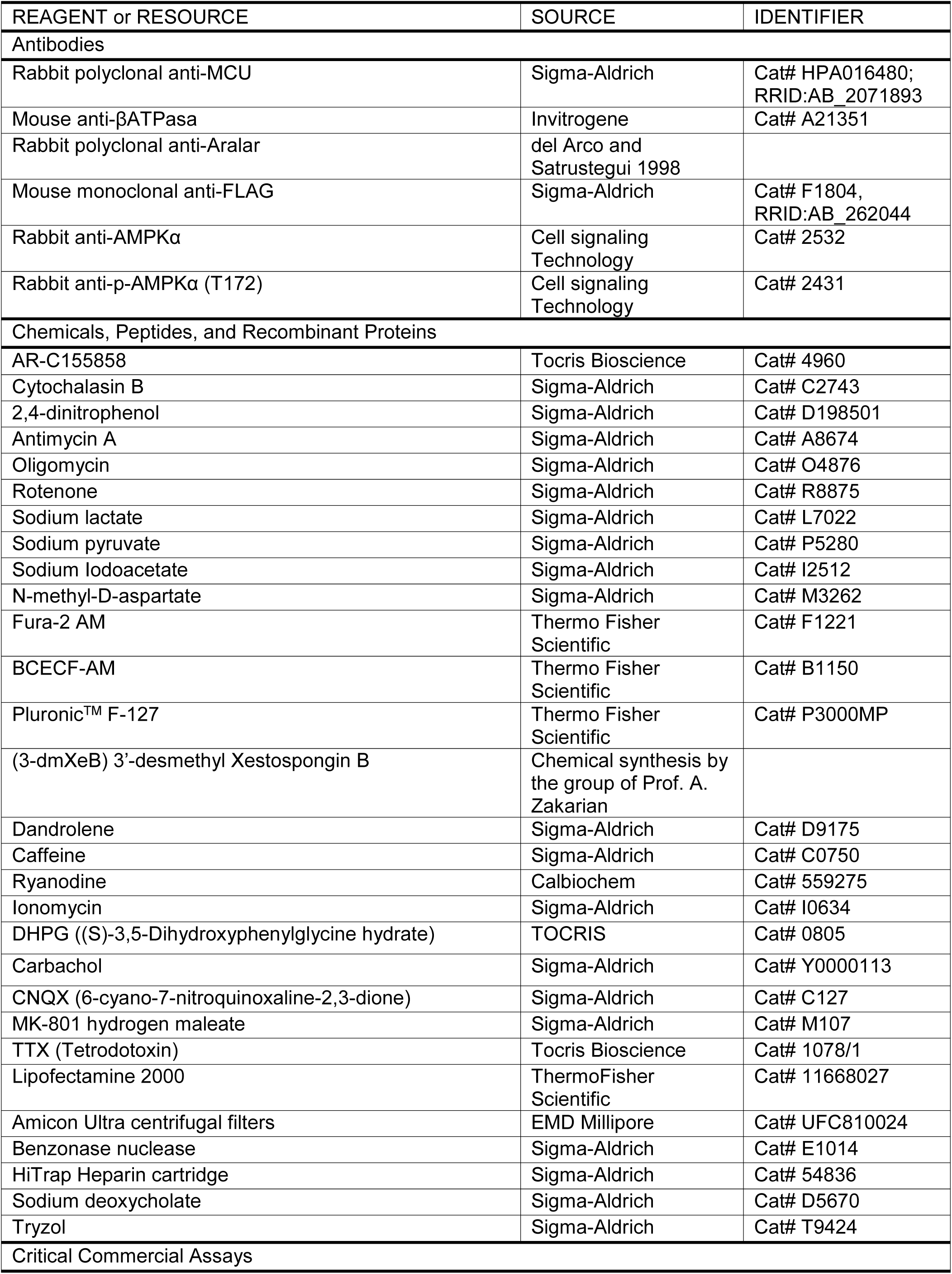
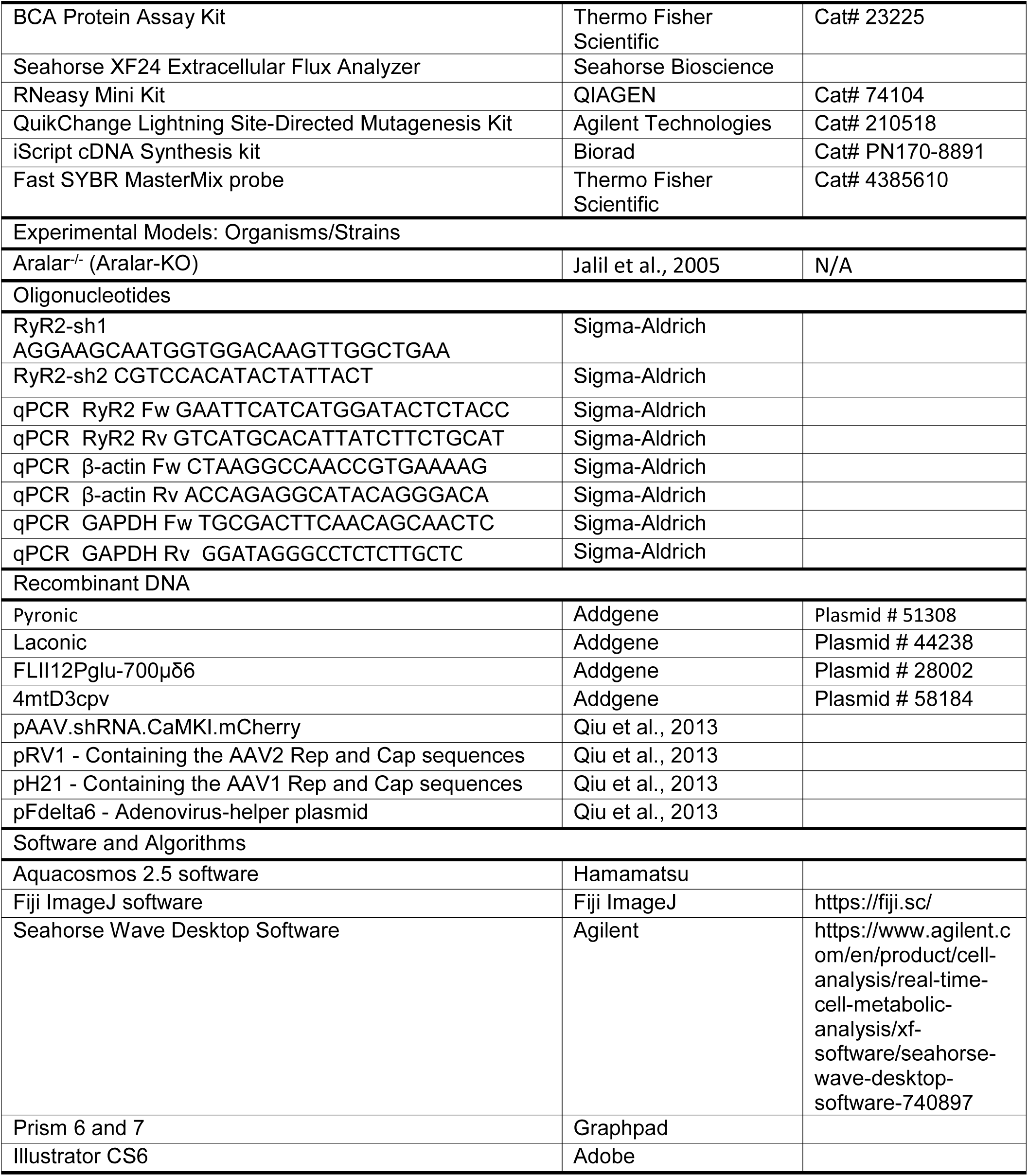

## Methods

### Animals

Mice with targeted disruption of *Aralar/AGC1/Slc25a12* (*Aralar*-KO) were described before (Jalil et al., 2005). *Aralar*-KO mice on a mixed SVJ129xC57BL/6 background were used. WT and *Aralar*-KO E16 embryos for preparing neuronal cultures were obtained by breeding of heterozygous *Aralar*-KO animals.

All animal procedures were performed in compliance with animal protocols approved by institutional ethical committee (Center of Molecular Biology Severo Ochoa) and Autónoma University (CEEA-CBMSO-23/159), and in accordance with Spanish regulations (BOE 67/8509-12, 1988) and European regulations (EU directive 86/609, EU decree 2001-486), reporting followed the ARRIVE Guidelines. All efforts were made to minimize the number of animals used and their suffering.

### Drugs

Cells were treated as indicated with cytochalasin B, 2,4-dinitrophenol (DNP), antimycin A (A), oligomycin (Olig), Rotenone (R), Caffeine, Ionomycin, Carbachol, Tetrodotoxin (TTX), dandrolene, N-methyl-D-aspartate (NMDA), MK-801 and 6-cyano-7-nitroquinoxaline-2,3-dione (CNQX) all purchased from Sigma-Aldrich, also with AR-C155858 and (S)-3,5-Dihydroxyphenylglycine hydrate (DHPG) from Tocris Bioscience and ryanodine from Calbiochem. Sodium lactate, sodium pyruvate and sodium Iodoacetate were from Sigma-Aldrich and used in the conditions indicated. 3-desmethyl Xestospongin B (3-dmXeB) was produced by chemical synthesis and dissolved in methanol. For *in vitro* experiments, compounds dissolved in DMSO or ethanol, both solvents were never higher than 0.001%. Corresponding vehicle solutions were used in control experiments. Doses and concentrations of the different drugs were chosen on the basis of previous published data or preliminary experiments.

### Primary neuronal culture

Neuronal cultures were obtained from E15-E16 mouse embryos, as described (Ruiz et al., 1998; Ramos et al., 2003; Pardo et al., 2006). Cerebral cortices were enzymatically dissociated in PBS containing 1% bovine serum albumin (BSA), 0.4 mg/ml papain (Roche), and 6 mM glucose.

Dissociated cells were plated on poly-L-lysine and laminin-coated glass coverslips in medium containing 20% horse serum. After 3 h, medium was completely replaced by serum-free Neurobasal medium (NB) supplemented with 2% B27, 1% glutamax (all from Gibco, Invitrogen) and 100 mg/ml penicillin-streptomycin. Neurons were maintained in these conditions until day *in vitro* (DIV) 8-9-10 for experimental analysis. Neurons represented more than 80% of the total cell population (Pardo et al., 2006).

### Constructs

For *Mcu* knockdown (KD) using shRNA, we used rAAV vectors with a U6 promoter driving shRNA expression that co-express also mCherry under control of CaMKII promoter to identify infected neurons, pAAV.shRNA.CaMKI.mCherry, kindly provided by Hilmar Bading and Giles E. Hardingham (Qiu et al., 2013). The Mcu hairpin and the scrambled one, used as control, have been described previously (Qiu et al., 2013). To generate shRNA *RyR2* constructs two target sequences, sh*RyR2*-1 and sh*RyR*2-2, used previously (More et al., 2018; Mu et al., 2014, respectively) were selected. RyR2-sh1, targeting sequences 5′-AGGAAGCAATGGTGGACAAGTTGGCTGAA-3′; and RyR2-sh2, 5 ‘-CGTCCACATACTATTACTC-3 ‘; were annealed and subcloned into the pAAV.shRNA.CaMKI.mCherry vector using standard molecular cloning techniques as described previously (Rueda et al., 2015). All constructs were verified by sequencing.

### rAAVs obtention

Viral particles (chimeric rAAVs containing the capsid proteins of both AAV1 and AAV2 at equal ratios) were produced for *Scr, RyR2*-KD and *Mcu*-KD. rAAV were purified and titrated, as described previously (McClure et al., 2011). Briefly, human HEK293 cells were co-transfected with the pAAV.shRNA.CaMKI.mCherry plasmid, the serotype-specific AAV helper plasmids, pH 21 and pRV-1, encoding rep and cap genes of AAV1 and AAV2, respectively, and the adenovirus helper plasmid (pFΔ6) by standard CaPO_4_ transfection. Cells were harvested 48 h after transfection, treated with benzonase (Sigma-Aldrich) to reduce the viscosity, and cell debris were cleared by centrifugation for 15 min. The rAAV particles were purified using HiTrap heparin affinity columns (Sigma-Aldrich) and concentrated using Amicon Ultra centrifugal filters (EMD Millipore) and solutions were pooled and sterile filtered. Primary cortical neurons were infected with 10^7^ rAAV particles/μL at DIV 3. Infection efficiencies were determined at DIV 9 by analyzing the fluorescence of mCherry, they ranged from 80 to 90% of the viable neurons. In infected neurons, oxygen consumption, calcium and fluorescence measurements were performed at DIV 9-10.

### Transfections

Primary cortical neurons, seeded on chambered coverglass 4-well plates (Lab-Tek) at 1.5 × 10^5^ cells/well density, were transfected at DIV 7-8 using calcium phosphate protocol with 1 µg of plasmids coding fluorescent sensors for the measurement of cytosolic glucose [FLII^12^Pglu-700μδ6; Addgene #28002 (Takanaga et al., 2008)], pyruvate [Pyronic; Addgene #51308 (San Martin et al., 2014)], lactate [Laconic; Addgene #44238 (San Martin et al., 2013)], and for mitochondrial Ca^2+^ determinations [4mt-D3cpv; Addgene #58184 (Palmer et al., 2006)]. For the transfection of rAAVs-transduced cells, neurons were seeded at higher density (2 × 10^5^ cells/well), and transfections with fluorescent sensors were performed using Lipofectamine 2000 (Invitrogen) following supplied instructions. FRET measurements were usually performed 24 or 48 h after transfection.

### Mitochondrial respiration measurements

Cellular oxygen consumption rate (OCR) was measured using Seahorse XF24 Extracellular Flux Analyzer (Agilent) as previously described (Llorente-Folch et al., 2013; Rueda et al., 2015; Juaristi et al., 2019a). At DIV 9-10, cells were equilibrated with bicarbonate-free low-buffered Dulbecco modified Eagle ‘s minimal essential medium (DMEM; without pyruvate, L-lactate, glucose, glutamine, or calcium) supplemented with 2.5 mM glucose and 2 mM CaCl_2_ for 1 h before XF assay. OCR was determined under basal conditions and after the sequential injections of medium or NMDA at the concentrations indicated and 6 μM oligomycin, 0.5 mM 2,4-dinitrophenol (DNP), and 1 μM rotenone/1 μM antimycin. Values of basal, ATP-linked, non-mitochondrial and mitochondrial uncoupled respiration (MUR) were determined (Qian and Van Houten, 2010; Brand and Nicholls, 2011). Protein from each well was extracted with 0.1% NP-40 PBS solution and quantified with BCA protein assay kit (Thermo Fisher), and data were normalized from protein concentration. Non-mitochondrial OCR was subtracted to OCR values and normalized from basal values. OCR/ECAR ratio values were calculated under basal conditions of OCR and extracellular acidification rate (ECAR) for each well.

### Cytosolic calcium and pH imaging

Primary cortical neurons were seeded on glass coverslips on 24-well plates at 2.0 × 10^5^, or 1.5 × 10^5^ for Carbachol stimulations, cells/well. At DIV 8-9, neurons were incubated during 30 min at 37°C in Hepes-buffered control salted solution (HCSS; 137 mM NaCl, 1.25 mM MgSO_4_, 10 mM HEPES, 3 mM KCl, 2 mM NaHCO_3_ and 1% BSA, pH 7.4) supplemented with 5 μM Fura-2 AM (Thermo Fisher), 50 μM pluronic acid F-127 (Thermo Fisher) and 2.5 mM glucose without calcium (plus 100 μM EGTA) for Ca^2+^ imaging, and with 0.12 μM BCECF-AM (Thermo Fisher), 50 μM pluronic acid F-127, 2.5 mM glucose and 2 mM CaCl_2_ for pH imaging. After loading, neurons were washed in HCSS with 2.5 mM glucose and 2 mM CaCl_2_ during 20 min at 37°C and then imaging was performed at 37°C as described (Ruiz et al., 1998; Juaristi et al., 2019a) with a Neofluar 40X/0.75 NA objective in an Axiovert 75M microscope (Zeiss). Acute additions were made as a bolus at the times indicated. Image acquisition was performed with Aquacosmos 2.5 software (Hamamatsu).

### Pyronic, laconic, FLII^12^Pglu-700μδ6 experiments

For pyronic, laconic, FLII^12^Pglu-700μδ6 experiments, cells were pre-incubated 1 h in HCSS (+Ca^2+^) at 37°C; for Ca^2+^-free assays, cells were washed once in HCSS (-Ca^2+^) prior imaging. Experiments were performed in HCSS supplemented with 2.5 mM glucose and 2 mM CaCl_2_ or 100 μM EGTA and the additions indicated. In pyronic and laconic experiments, 2 mM pyruvate or lactate were added at the end of the experiment to verify that the probe was not saturated. Neurons were excited at 430 nm (CFP), and emission was collected with a dual pass dichroic filter CFP-YFP (440-500 nm and 510-610 nm). Exposure time was 200 ms. Images were collected every 5 seconds using a filter wheel Lambda 10-3, Sutter Instruments (Chroma) and recorded by a Hamamatsu C9100-02 camera mounted on an Axiovert 200M inverted microscope equipped with a 40X /1.3 oil Plan-Neofluar objective.

Regions of interest (ROIs) containing somas were selected on each cell and analysed using MetaMorph (Universal Imaging) and ImageJ (NIH, https://fiji.sc/) software. Background was subtracted from YFP and CFP images and subsequently, the ratio between CFP and YFP emissions calculated. Rates of fluorescence increase in response to NMDA stimulation were calculated as the lineal increase in fluorescence ratio (ΔF/min) during the first 30 seconds after addition. Maximal fluorescence ratio increase (ΔF max) and fluorescence ratio at the end of the recording (ΔF endpoint) after NMDA stimulation were calculated.

### Adenine nucleotide quantification

Adenine nucleotides (AdN) total levels were determined by HPLC (de Korte et al., 1995). Neuronal cultures were treated with 660 mM HClO_4_ and 10 mM theophylline immediately frozen and kept them over-night at -80°C. Neurons were scrapped and homogenized on ice, then centrifuged at 16000 g 15 min 4°C. The supernatants were neutralized using 2.8 M K_2_CO_3_, kept on ice 10 min, and then at -80°C for at least 2 h to allow precipitation of HClO_4_. Extracts were centrifuged and 50 µl of the supernatants were injected for AdN determination. To determine the identity of each peak and quantify the amount of the species of interest, standard curves were constructed by plotting peak heights versus known concentrations of ATP, ADP, and AMP.

### Quantitative RT-PCR

RNA from primary cortical neurons was isolated using Trizol reagent (TRIzol; Sigma-Aldrich) following manufacturer ‘s instructions. Retrotranscription (RT) reactions were performed using the iScript cDNA Synthesis kit (Biorad PN170-8891) following manufacturer’s instructions. Thermal conditions consisted of the following steps: 5’x 25°C, 20’x 46°C and 1’x 95°C. cDNA was amplified with Fast SYBR MasterMix probe (Thermo Fisher Scientific) in the ABI Prism 7900HT sequence detection system (Thermo Fisher Scientific) at the Genomics and Massive Sequencing Facility (CBMSO–UAM). The primers used for amplifying the target genes were: mouse RyR2 (5 ‘-GAATTCATCATGGATACTCTACC-3 ‘; 5 ‘-GTCATGCACATTATCTTCTGCAT-3 ‘). Mouse β-actin (5 ‘-CTAAGGCCAACCGTGAAAAG-3 ‘; 5 ‘-ACCAGAGGCATACAGGGACA-3 ‘) and mouse GAPDH (5 ‘-TGCGACTTCAACAGCAACTC-3 ‘; 5 ‘-GGATAGGGCCTCTCTTGCTC-3 ‘) were used as housekeeping genes. Valid Prime expression (Tataa Biocenter, A106S10) was checked in all the samples and necessary corrections were done as described in Laurell et al. (2012). The relative expression of genes was calculated using the 2^-ΔΔCt^ Livak comparative method (Schmittgen and Livak, 2008).

### Western blotting and antibodies

To examine protein levels, total protein extracts were prepared in RIPA buffer (125 mM NaCl, 25 mM Tris-Cl pH 7.4, 1 mM EGTA, 1% Triton X-100, 0.5% sodium deoxycholate, 0.1% SDS and Complete EDTA-free protease inhibitor (Roche). 40 µg of total proteins were loaded and were separated by SDS-PAGE electrophoresis, and transferred onto nitrocellulose membranes by wet electrophoretic transfer. Blots were blocked 1 h at RT with 5% non-fat dry milk in TBS-tween (0.5 M Tris, 1.5 M NaCl, 0.01% Tween) solution and incubated at 4°C with primary antibodies. For phospho-antibodies, 5% BSA in TBS-tween was used as blocking solution. Secondary antibodies were incubated 1 h at RT. The following antibodies were used: anti-MCU (1:1,000, Sigma-Aldrich), anti-AMPKα (1:1,000, Cell Signaling Technology), anti-p-AMPK T172 (1:1,000, Cell Signaling Technology), anti-β-F_1_ATPase (1:5,000, kindly provided by Jose M. Cuezva). Secondary HRP-conjugated antibodies were purchased from Bio-Rad and used at 1:5,000 dilution. The signal was detected by chemiluminescence with detection solution (75 mM Tris pH 8.8, 2.5 mM luminol, 0.40 mM p-coumaric acid, 0.01% H_2_O_2_). The quantification of the bands was performed with ImageJ (NIH) software.

### Quantification and statistical analysis

Statistical details of experiments can be found in the figure legends. All results are representative of at least 3 independent experiments unless otherwise specified and are presented as mean ± SEM. Significance was calculated by ANOVA (one-way or two-way), *post hoc* Bonferroni test or T-test. All statistical tests were plotted either with Sigma Plot 11.0 software or GraphPad Prism version 6 or 7 software.

---

**Figure 1 supplementary 1.**
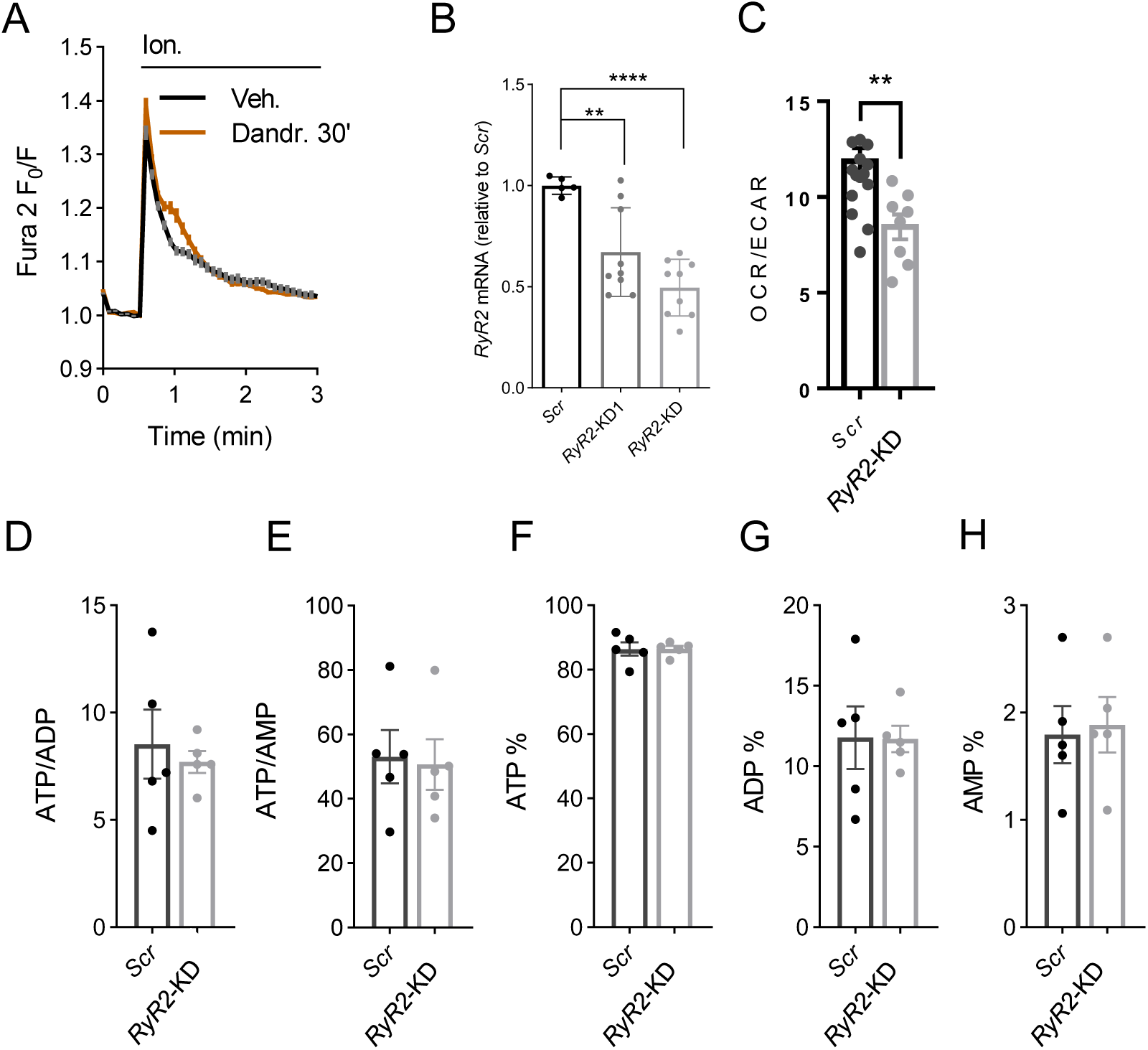
Adenine nucleotide content in *RyR2*-KD neurons. A: No changes in 5 µM inomycin (Ion.) produced ER calcium discharge between control neurons and neurons incubated during 30 min with 10 µM dandrolene (Dandr.). B: qPCR analysis of the cDNA obtained from neuronal cultures transduced with rAAV containing two different *RyR2*-directed (*RyR2*-KD1 or *RyR2*-KD) or Scrambled shRNA (Scr) using specific oligos; all values shown are relative to the expression levels of RyR2 mRNA in *Scr* neurons. Mean ± SEM (bars), 5-9 wells (dots) from two independent experiments; one-way ANOVA, \*\**p* ≤ 0.01, \*\*\*\**p* ≤ 0.0001 post hoc Bonferroni test. B: RyR2 silencing decreases basal OCR/ECAR ratio values. Mean ± SEM (bars), n = 17-8, from 3 independent experiments; two -tailed T-test, \*\**p*-value = 0.0041. D-H: Adenine nucleotide content, measured by HPLC in neurons transduced with rAAV containing *RyR2*-directed (*RyR*-KD) or Scrambled shRNA (*Scr*). Mean ± SEM (bars), 5 independent experiments (dots).

**Figure 3 Supplementary 1.**
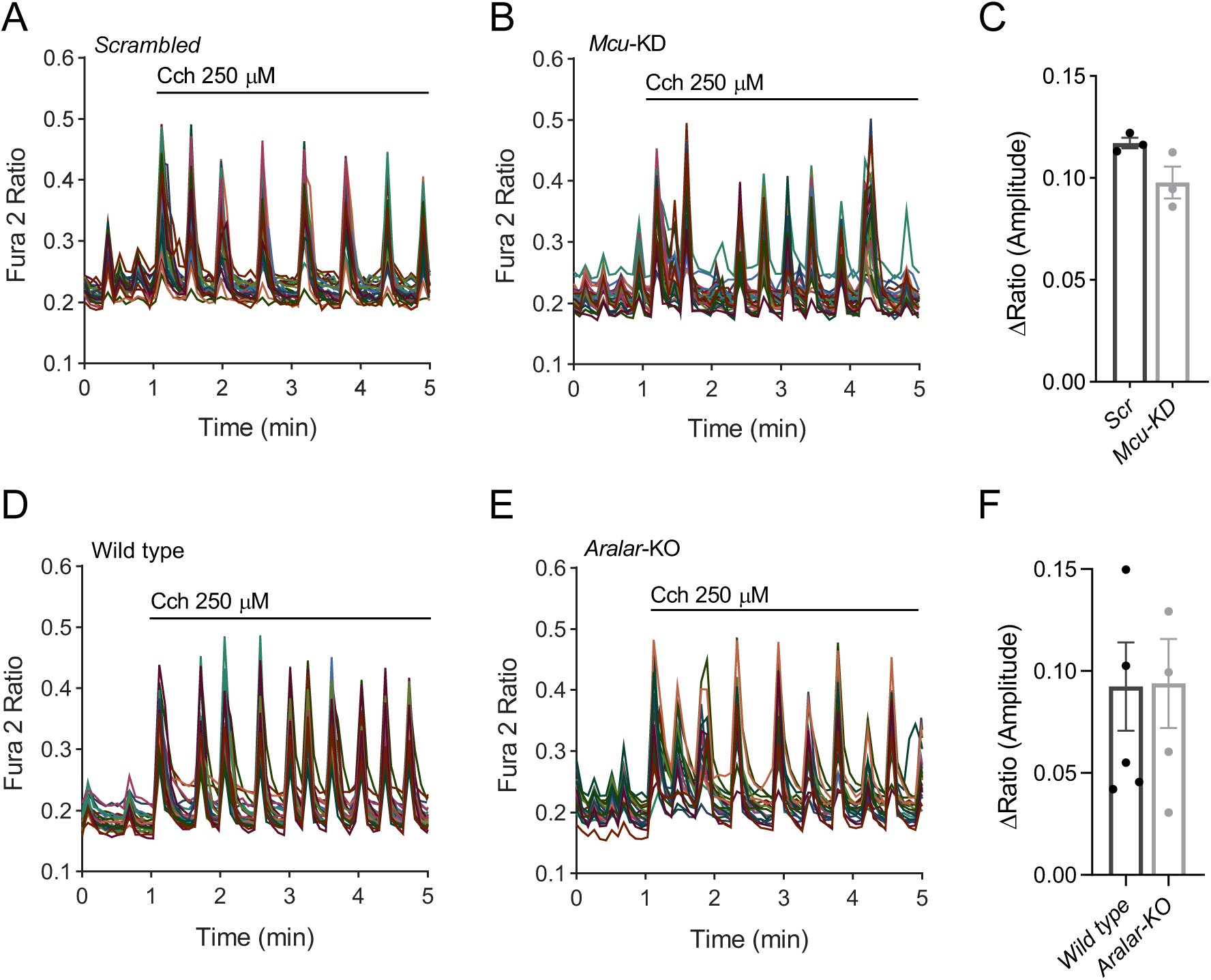
Cch induced cytosolic Ca^2+^ signals. A-B: Fura-2AM [Ca^2+^]i signals in *Scr* (A) or *Mcu*-KD (B) neurons in HCSS medium containing 2 mM CaCl_2_ and 2.5 mM glucose upon addition of 250 μM Cch where indicated. Representative experiments, each trace corresponds to a single neuron from the same recording field. C: Quantification of peak amplitude as ΔRatio (F_340_/F_380_) ± SEM. Data were obtained from 3 independent experiments. D-E: Fura-2AM [Ca^2+^]i signals in WT (D) and *Aralar*-KO (D) neurons in HCSS medium containing 2 mM CaCl_2_ and 2.5 mM glucose upon addition of 250 μM Cch where indicated. Representative experiments, each trace corresponds to a single neuron from the same recording field. F: Quantification of peak amplitude as ΔRatio (F_340_/F_380_) ± SEM. Data were obtained from 5 independent experiments.

**Figure 4 supplementary 1.**
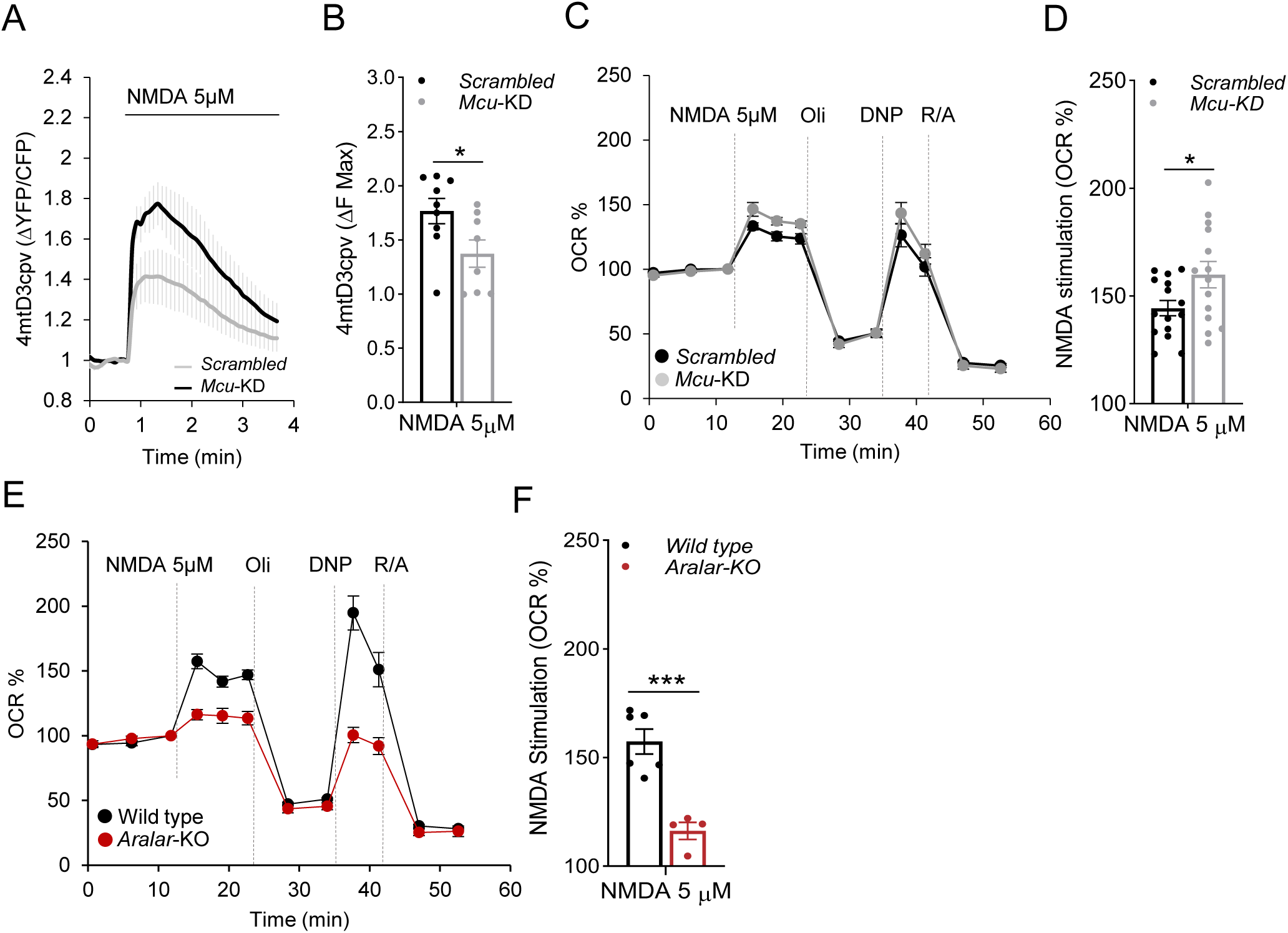
Changes in mitochondrial Ca^2+^ and OCR upon stimulation with 5 μM NMDA. A: Changes in mitochondrial Ca^2+^ (Ca^2+^-mit) in *Scrambled* (*Scr*) and *Mcu*-silenced (*Mcu*-KD) neurons transfected with 4mtD3cpv probe, after 5 μM NMDA stimulation in 2.5 mM glucose and 2 mM Ca^2+^ HCSS; B: Quantification of maximum Ca^2+^-mit increment (ΔF max) after NMDA stimulation. Data are mean ± SEM (bars), *n* = 8-9 (dots) from three independent experiments; T-test \**p*-value = 0.0365. C: 5 μM NMDA-stimulated respiration in *Scrambled* and *Mcu*-KD neurons maintained in 2.5 mM glucose and 2 mM Ca^2+^ DMEM, expressed as percentage of basal values (OCR %); D: Quantification of NMDA-stimulated OCR (%); mean ± SEM (bars), *n* = 14-15 (dots) from 5 independent experiments; T-test \**p*-value = 0.0339. E: 5 μM NMDA-stimulated respiration in wild type and *Aralar*-KO neurons maintained in 2.5 mM glucose and 2 mM Ca^2+^ DMEM, expressed as percentage of basal values (OCR %); F: Quantification of NMDA-stimulated OCR (%); mean ± SEM (bars), *n* = 4-6 (dots) from 2 independent experiments; T-test \*\*\**p*-value = 0.0007. Mitochondrial function was determined through sequential addition of metabolic inhibitors: 6 μM Olig, 0.5 mM DNP and 1 μM/1 μM A/R at the indicated time points.

**Figure 4 supplementary 2.**
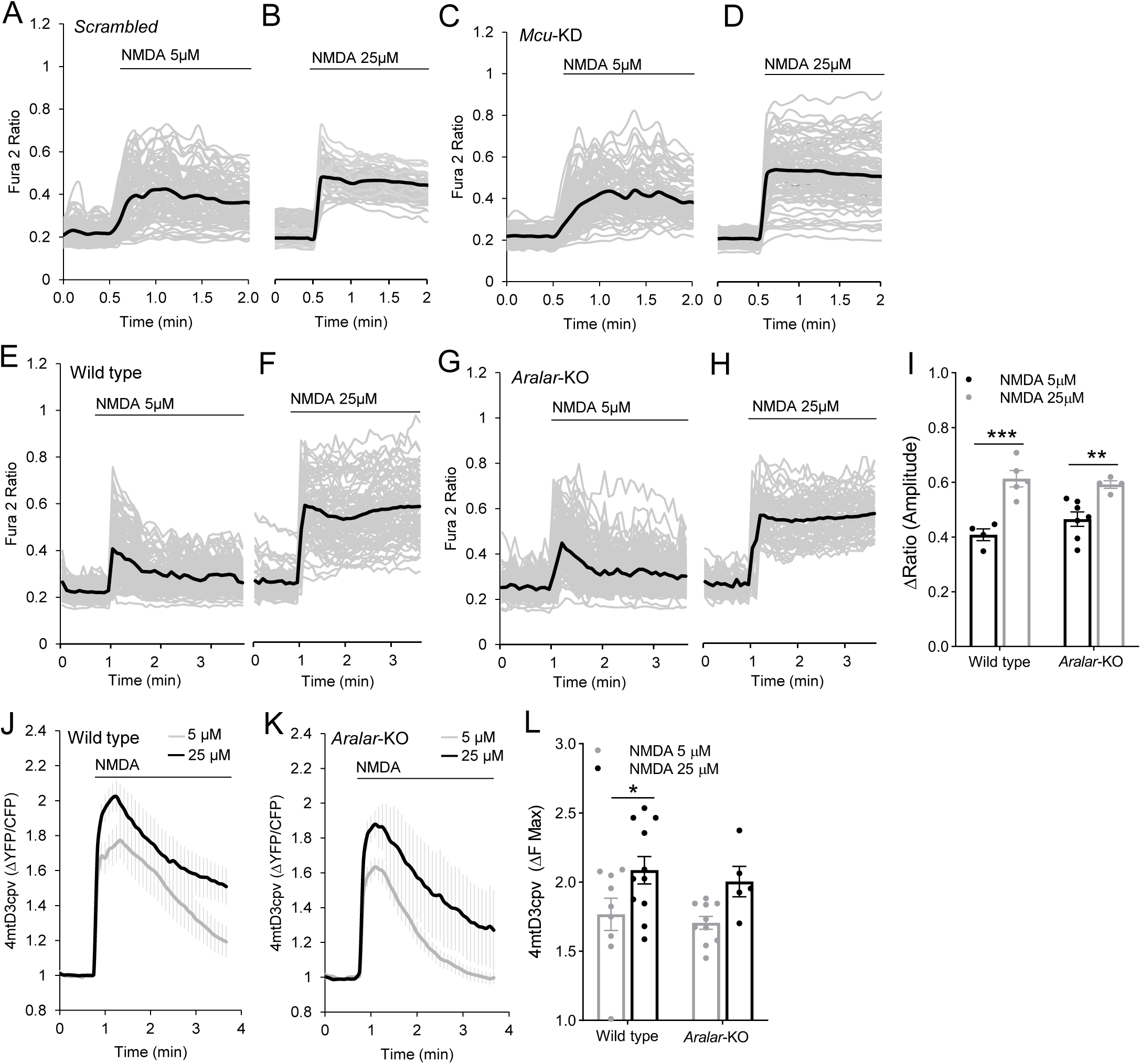
Cytosolic and mitochondrial Ca^2+^ signals in *Scr*, *Mcu*-KD, Wild type and *Aralar*-KO neurons. A-D: Changes in cytosolic Ca^2+^ (Ca^2+^-cyt) after 5 (A, C) and 25 μM NMDA (B, D) stimulation in Scr (A, B) or Mcu-KD (C, D) neurons. E-I: Changes in cytosolic Ca^2+^ (Ca^2+^-cyt), after 5 (E, G) and 25 μM (F, H) NMDA in wild type (E, I) and *Aralar*-KO (G, H) Fura2-AM loaded neurons. Individual cell recordings (grey) and average (black) are shown. I: Quantification of peak amplitude as ΔRatio (F_340_/F_380_) ± SEM (bars) comparing 5 and 25 μM NMDA stimulus; two-way ANOVA, \*\**p* ≤ 0.01,\*\*\**p* ≤ 0.001, post hoc Bonferroni test. Recordings from at least 50 cells/condition from 4-7 wells (dots) and two independent experiments. J-L: Changes in mitochondrial Ca^2+^ (Ca^2+^-mit), in wild type (J) and *Aralar*-KO neurons (K) transfected with 4mtD3cpv probe, after 5 and 25 μM NMDA stimulation; L: maximum Ca^2+^-mit increment (ΔF max) after NMDA stimulation. Mean ± SEM (bars), 5-10 cells (dots) from two independent culture; two-way ANOVA, \**p* ≤ 0.05, post hoc Bonferroni test. All cells were maintained in 2.5 mM glucose and 2 mM Ca^2+^ HCSS.

**Figure 5 supplementary 1.**
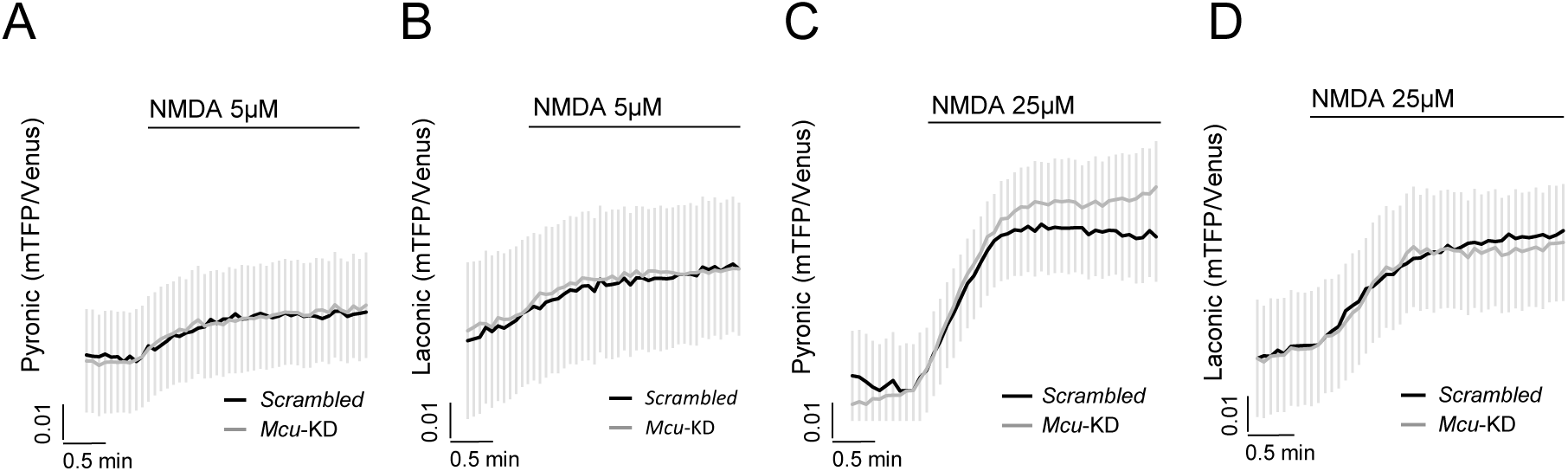
Effects of NMDA stimulation on cytosolic pyruvate and lactate production in Mcu-silenced neurons. Cytosolic pyruvate (Pyr) (A, C) and lactate (Lac) (B, D) levels of Scr and *Mcu*-KD silenced neurons transfected with Pyronic and Laconic probes, respectively, upon neuronal stimulation with 5 μM (A, B) or 25 μM NMDA (C, D). Neurons were incubated in 2.5 mM glucose and 2 mM Ca^2+^ HCSS. Data are mean ± SEM, *n* = 7-15 neurons per condition from 2 independent cultures.

